# NAMPT-derived NAD^+^ fuels PARP1 to promote skin inflammation through parthanatos

**DOI:** 10.1101/2021.02.19.431942

**Authors:** Francisco J. Martínez-Morcillo, Joaquín Cantón-Sandoval, Francisco J. Martínez-Navarro, Isabel Cabas, Idoya Martínez-Vicente, Joy Armistead, Julia Hatzold, Azucena López-Muñoz, Teresa Martínez-Menchón, Raúl Corbalán-Vélez, Jesús Lacal, Matthias Hammerschmidt, José C. García-Borrón, Alfonsa García-Ayala, María L. Cayuela, Ana B. Pérez-Oliva, Diana García-Moreno, Victoriano Mulero

## Abstract

Several studies have revealed a correlation between chronic inflammation and NAD^+^ metabolism but the precise mechanism involved is unknown. Here we report that the genetic and pharmacological inhibition of nicotinamide phosphoribosyltransferase (Nampt), the rate-limiting enzyme in the salvage pathway of NAD^+^ biosynthesis, reduced oxidative stress, inflammation, and keratinocyte DNA damage, hyperproliferation and cell death in zebrafish models of chronic skin inflammation, while all these effects were reversed by NAD^+^ supplementation. Similarly, genetic and pharmacological inhibition of poly ADP-ribose (PAR) polymerase 1 (Parp1), overexpression of PAR glycohydrolase, inhibition of apoptosis-inducing factor 1, inhibition of NADPH oxidases and reactive oxygen species (ROS) scavenging, all phenocopied the effects of Nampt inhibition. Pharmacological inhibition of NADPH oxidases/NAMPT/PARP/AIFM1 axis decreased expression of pathology-associated genes in human organotypic 3D skin models of psoriasis. Consistently, an aberrant induction of both NAMPT amounts and PARP activity was observed in lesional skin from psoriasis patients. In conclusion, hyperactivation of PARP1 in response to ROS-induced DNA damage, fueled by NAMPT-derived NAD^+^, mediates skin inflammation through parthanatos cell death.

**Highlights:**

- NAMPT inhibition alleviates inflammation in zebrafish and human epidermis organoid models of psoriasis.
- NADPH oxidase-derived ROS mediates keratinocyte DNA damage and Parp1 overactivation.
- Inhibition of parthanatos cell death phenocopies the effects of NAMPT inhibition in zebrafish and human psoriasis models.
- NAMPT and PAR metabolism is altered in psoriasis patients.

## Introduction

Psoriasis is a non-contagious chronic inflammatory skin disease with global prevalence of 0.1-3 % (Guttman-Yassky and Krueger, 2017). Despite being a relapsing and disabling disease that affects both physical and mental health, is not usually life threatening. However, the cytokines and chemokines produced in the lesion may reach the blood and consequently cause comorbidities (Furue and Kadono, 2017). Although the etiology is still undetermined, external agents trigger inflammation in genetically predisposed epithelium (Greb et al., 2016; Weidinger et al., 2018). Cytokines released by keratinocytes stimulate dendritic/Langerhans cells that drive specific Th17 cell immune response and additional cytokines and chemokines close the inflammatory feed-back loop that result in the skin lesion (Dainichi et al., 2018).

Nicotinamide adenine dinucleotide (NAD^+^) is the most important hydrogen carrier in redox reactions in the cell, participating in vital cellular processes such as mitochondrial function and metabolism, immune response, inflammation and DNA repair, among others (Rajman et al., 2018). NAD^+^ levels are tightly regulated by Preiss-Handler, *de novo* and salvage pathways (Canto et al., 2015). Different tissues preferentially employ a distinct pathway regarding available precursors. The most important NAD^+^ precursor is dietary niacin (also known as vitamin B3), consisting of nicotinamide (NAM), nicotinic acid (NA) and NAM riboside (NR) (Houtkooper et al., 2010). NAM is also the product of NAD^+^-consuming enzymes, that is why most mammalian tissues rely on NAM to maintain the NAD^+^ pool, via the NAD^+^ salvage pathway (Canto et al., 2015; Rajman et al., 2018). The rate-limiting enzyme in the NAD^+^ salvage pathway is NAM phosphoribosyltransferase (NAMPT) that converts NAM into nicotinamide mononucleotide (NMN). After that, NMN adenylyltransferases (NMNAT 1-3) transform NMN into NAD^+^ (Houtkooper et al., 2010). NAMPT has been associated with oxidative stress and inflammation (Garten et al., 2015), being identified as a universal biomarker of chronic inflammation, including psoriasis (Mesko et al., 2010). FK-866 is a noncompetitive highly specific NAMPT pharmacological inhibitor that induces a progressive NAD^+^ depletion (Hasmann and Schemainda, 2003). FK-866 has demonstrated anti-inflammatory effects in different experimental settings, including murine models of colitis and collagen-induced arthritis (Busso et al., 2008; Gerner et al., 2018).

Several enzymes depend on NAD^+^ to accomplish their biological functions. Poly(ADP-Ribose) (PAR) polymerases (PARPs) are major NAD^+^-consuming enzymes which transfer ADP-ribose molecules (linear or branching PAR) to proteins or itself (auto-PARylation) (Qi et al., 2019). PARPs are implicated in DNA repair and chromatin organization, gene transcription, inflammation and cell death or stress responses, among others (Canto et al., 2015; Qi et al., 2019). However, the main PARP biological function is to orchestrate the spatio-temporal repair of DNA damage; that is why PARP1 is predominantly localized in the nucleus (Canto et al., 2015), being responsible for approximately 90 % of PAR biosynthesis (Qi et al., 2019). PARP1-3 are recruited and activated upon single- and double-strand DNA breaks (ssBs and dsBs) (Qi et al., 2019). Several PARP inhibitors have been developed, such as olaparib, niraparib, rucaparib, talazoparib and veliparib (Kukolj et al., 2017; Nesic et al., 2018). Once PARP1 is recruited to a ssB, the inhibitors prevent PARP enzymatic activity, entrapping and accumulating inactive PARP on DNA and triggering the collapse of replication forks resulting in dsB generation during replication (Kukolj et al., 2017).

Under physiological conditions, DNA damage provoked by cellular metabolism is successfully handled by PARP1. However, alkylating DNA damage, oxidative stress, hypoxia, hypoglycemia or activation of inflammatory pathways can trigger PARP1 hyperactivation. Excessive PARylation depletes cellular NAD^+^ and ATP stores, although it does not directly imply cell death. However, the accumulation of PAR polymers and PARylated proteins reach the mitochondria causing depolarization of the membrane potential and apoptosis-inducing factor (AIFM1) release into the cytosol (Fatokun et al., 2014; Galluzzi et al., 2018). AIFM1 then recruits macrophage migration inhibitory factor (MIF) to the nucleus where AIFM1-MIF nuclease activity executes a large-scale DNA fragmentation resulting in a cell death pathway known as parthanatos (Wang et al., 2016).

In this work, we report a critical role played by NAD^+^ and PAR metabolism in skin oxidative stress and inflammation. We found that hyperactivation of PARP1 in response to ROS-induced DNA damage, and fuelled by NAMPT-derived NAD^+^, mediates inflammation through parthanatos cell death in preclinical zebrafish and human epidermis organoid models of psoriasis. Consistent with this, clinical data support the alteration of NAD^+^ and PAR metabolism in psoriasis, pointing to NAMPT and PARP1 as novel therapeutic targets to treat skin inflammatory disorders.

## Results

### NAD^+^ metabolites regulate skin oxidative stress and inflammation

In order to determine if NAD^+^ metabolism has any role in the regulation of skin inflammation, we decided to perform functional experiments in the transgenic zebrafish line *lyz:dsRED,* which labels neutrophils. Manually dechorionated *lyz:dsRED* larvae were treated by bath immersion with different concentrations of NAD^+^ from 24 hpf to 72 hpf (Figure 1A). Incubation with 1 mM NAD^+^ resulted in a statistically significant increased neutrophil dispersion from the caudal hematopoietic tissue (CHT) compared to vehicle (DMSO)- and 0.25 and 0.5 mM NAD^+^-treated larvae (Figure 1A-1C). Despite the altered pattern of neutrophil distribution, some of which were present in the skin, both the integrity of the skin and its morphology were not affected (Figure 1C).

**Figure 1.**
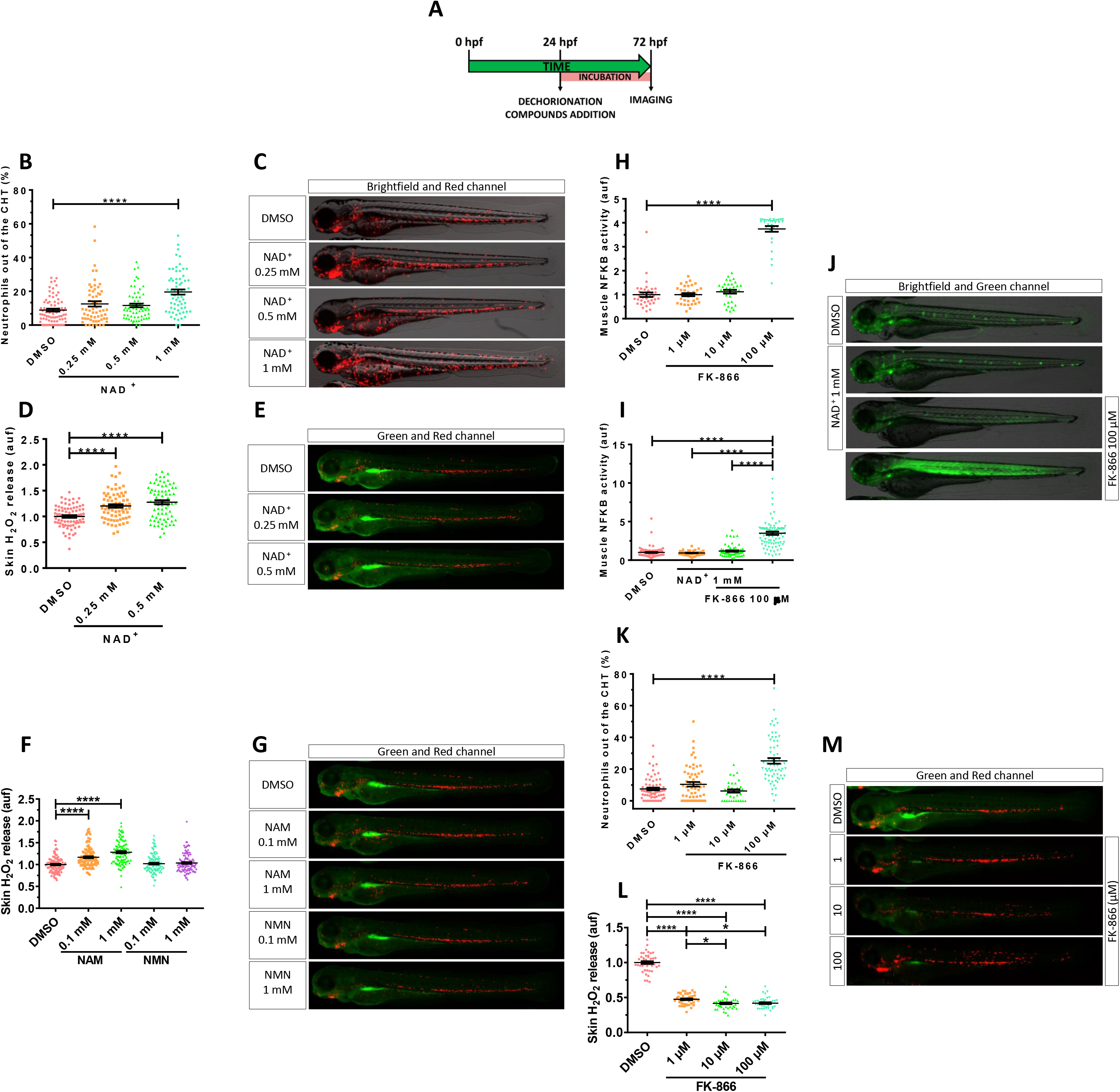
NAD^+^ metabolites regulate skin oxidative stress and inflammation. (A) Embryos of 24 hpf were manually dechorionated, treated for 48 hours with NAD^+^ metabolites or FK-866 by bath immersion and images obtained at 72 hpf. (B-M) Quantification of the percentage of neutrophils out of the CHT in embryos treated with NAD^+^ (0.25, 0.5 and 1 mM) (B). Representative merge images (brightfield and red channel) of *lyz:dsRED* zebrafish larvae of every group are shown (C). For H_2_O_2_ imaging, embryos were incubated in 50 µM of acetyl-pentafluorobenzene sulphonyl fluorescein solution for 1 hour. Quantification of fluorescence intensity for NAD^+^-mediated (D) and NAM-/NMN-mediated (F) induction of H_2_O_2_ in the zebrafish skin. Representative merge images (green and red channel) of *lyz:dsRED* zebrafish larvae of every group are shown (E, G). NFKB activity was determined by quantification of fluorescence intensity in embryos treated with increasing doses of FK-866 (1, 10 and 100 µM) (H). Additionally, the influence of NAD^+^ in the presence or absence of 100 µM FK-866 was assayed (I) Representative images (brightfield and green channel) of *nfkb:eGFP* zebrafish larvae of every group are shown (J). Neutrophil distribution in zebrafish embryos of 3 dpf treated with FK-866 (K), quantification of skin H_2_O_2_ production (L) and **r**epresentative merge images (green and red channel) of *lyz:dsRED* zebrafish larvae of every group are shown (M). Each dot represents one and the mean ± S.E.M. for each group is shown. P values were calculated using one-way ANOVA and Tukey multiple range test. *p≤0.05, ****p≤0.0001.

Given the role of H2O2 in driving neutrophil mobilization to acute (Niethammer et al., 2009) and chronic (Candel et al., 2014) insults, we used the H2O2 fluorescent probe acetyl-pentafluorobenzene sulphonyl fluorescein to know if this molecule was involved in the observed phenotype. NAD^+^ treatment was able to enhance H2O2 production in skin in a dose-dependent manner compared to the control group (Figure 1D, 1E). Similar results were obtained with NAM (Figures 1F, 1G), a well-known NAD^+^ booster (Rajman et al., 2018), while NMN precursor was unable to increase skin oxidative stress (Figures 1F, 1G). Nevertheless, no differences in neutrophil redistribution were observed (Figure 1G).

We next wondered if the depletion of cellular NAD^+^ stores by the well-characterized NAMPT inhibitor FK-866 (Hasmann and Schemainda, 2003) could also have an impact on skin oxidative stress and inflammation using the H2O2 probe and the transgenic line *nfkb:eGFP*, which accurately reports the activity of the master inflammation transcription factor NFKB (Kanther et al., 2011). FK-866 promoted a robust induction of NFKB activity in the muscle at the highest concentration used (100 µM) compared to control group (Figures 1H, 1J). Despite the NAD^+^ ability to induce skin oxidative stress when used at 1 mM, it was unable to activate NFKB transcriptional activity in any tissue. Importantly, NAD^+^ effectively restored muscle NFKB activity in FK-866-treated larvae (Figures 1I, 1J), confirming the specificity of the inhibitor.

The inflammatory effect of FK-866 in muscle was confirmed by the robust neutrophil infiltration (Figure 1K, 1M). Furthermore, H2O2 production by skin keratinocytes was almost abolished by 1 µM of FK-866 (Figure 1L, 1M). Collectively, these results suggest that not only NAD^+^ metabolite levels regulate oxidative stress in the skin and neutrophil infiltration, but also that low levels of NAD^+^ trigger muscle inflammation.

### Inhibition of Nampt alleviates oxidative stress and skin inflammation in a zebrafish model of psoriasis

The influence of NAD^+^ metabolism on skin oxidative stress and inflammation in wild type zebrafish, encouraged us to study its effect on the zebrafish psoriasis model with an hypomorphic mutation of *spint1a* (allele *hi2217*), which encodes the serine protease inhibitor, kunitz-type, 1a. Spint1a-deficient larvae showed increased H2O2 release in the skin compared with their wild type siblings (Figures 2A, 2B). As described above for wild type larvae, pharmacological inhibition of Nampt with FK-866 robustly decreased in a dose-dependent manner H2O2 production by Spint1a-deficient skin keratinocytes (Figures 2A, 2B).

**Figure 2.**
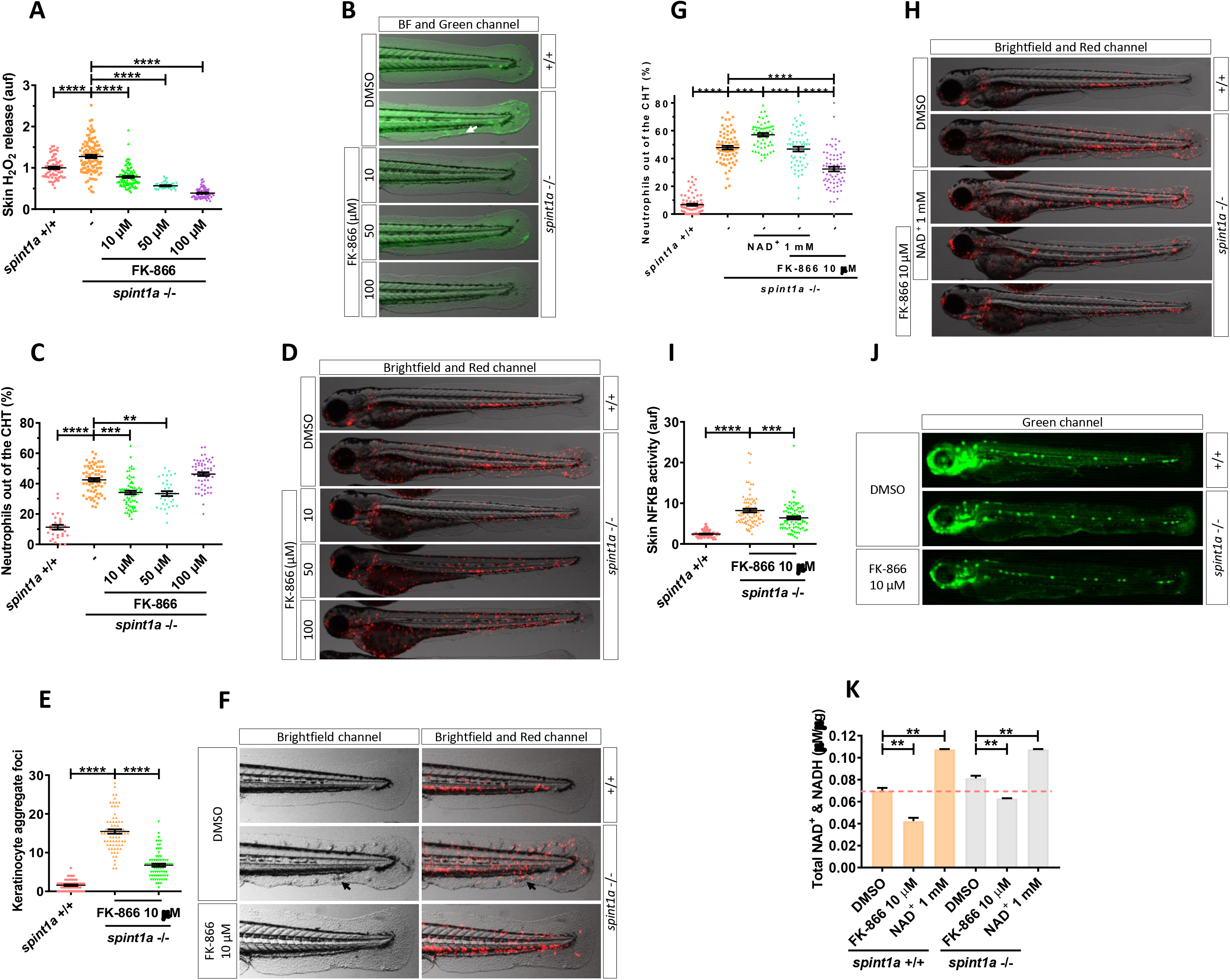
Pharmacological inhibition of Nampt alleviates stress and skin inflammation in Spint1a-deficient larvae. (A) Quantification of skin H_2_O_2_ production in wild type and Spint1a-deficient zebrafish larvae treated with FK-866 (0, 10, 50 and 100 µM). (B) Representative merge images (brightfield and green channel) of zebrafish larvae of every group are shown. The white arrow indicates a keratinocyte aggregate where H_2_O_2_ release was high. (C) Neutrophil distribution of wild type and Spint1a-deficient zebrafish larvae treated with increasing doses of FK-866. (D) Representative merge images (brightfield and red channel) of *lyz:dsRED* zebrafish larvae of every group are shown. (E) Number of keratinocyte aggregates in the tail skin of larvae treated with 10 µM FK-866 (F) Detailed representative merge images (brightfield and red channel) of the tail of wild type and Spint1a-deficient larvae treated with vehicle (DMSO) or 10 µM FK-866. Black arrows indicate keratinocyte aggregates and immune cell recruitment presents in non-treated *spint1a*-deficient skin. (G) Neutrophil distribution of zebrafish embryos treated with 1 mM NAD^+^ in the presence or absence of 10 µM FK-866. (H) Representative merge images (brightfield and red channels) of *lyz:dsRED* zebrafish larvae of every group are shown. (I) Quantification of fluorescence intensity of wild type and *spint1a* mutant embryos treated with 10 µM FK-866. (J) Representative images (green channel) of *NF-kB:eGFP* zebrafish larvae of every group are shown. (K) Wild type and Spint1a-deficient larvae of 72 hpf treated for 48 h with 10 µM FK-866 and 1 mM NAD^+^ were used for total NAD^+^ and NADH determination by ELISA. Each dot represents one individual and the mean ± S.E.M. for each group is also shown. P values were calculated using one-way ANOVA and Tukey multiple range test.**p≤0.01, ***p≤0.001, ****p≤0.0001.

Spint1a-deficient larvae show neutrophil infiltration in the skin (Carney et al., 2007; LeBert et al., 2015; Mathias et al., 2007). We observed that 40% of neutrophils were out of the CHT in the mutants compared to 10% in wild type larvae (Figures 2C, 2D). Mutant larvae treated with 10 or 50 µM FK-866 displayed a strong reduction of neutrophil dispersion (Figures 2C, 2D). However, 100 µM FK-866 induced muscle neutrophil infiltration, as observed in wild type animals (Figures 2C, 2D). Importantly, epithelial integrity and keratinocyte aggregate foci were almost completely rescued in Spint1a-deficient larvae treated with 10 µM FK-866 (Figures 2E, 2F) or with another FK-866 inhibitor (GMX1778) (Figure S1A, S1B). These results were further confirmed by genetic inhibition of the 2 zebrafish Nampt paralogues, namely Nampta and Namptb (Figure S1C-S1F). In addition, NAD^+^ supplementation exerted a negative effect on Spint1a-deficient skin. Thus, NAD^+^ aggravates skin morphology alterations and neutrophil infiltration compared with wild type animals (Figures 2G, 2H). In addition, NAD^+^ treatment neutralized the beneficial effects of FK-866 on the *spint1a* mutant, worsening skin alterations and neutrophil infiltration (Figures 2G, 2H). Consistently, the high NFKB transcriptional activity observed in the skin of Spint1a-deficient larvae was reduced by FK-866 (Figures 2I, 2J). As expected, FK-866 and NAD^+^ supplementation decreased and increased, respectively NAD^+^/NADH levels (Figure 2K). However, no statistically significant differences between NAD^+^/NADH levels in wild type and Spint1a-deficient larvae were found (Figure 2K). Collectively, these results indicate that Spint1a-deficient animals were more susceptible to NAD^+^ supplementation than their wild type siblings and that the beneficial effects of FK-866 on the skin were mediated by reducing skin NAD^+^ availability.

### NADPH oxidase-derived ROS promote skin inflammation in Spint1a-deficient larvae

The higher levels of ROS in the skin of Spint1a-deficient larvae, together with their drastic reduction by pharmacological inhibition of Nampt, led us to hypothesize that Nampt-derived NAD^+^ was fuelling NADPH oxidases. We therefore used N-acetylcysteine (NAC), which can scavenge ROS directly and replenish reduced glutathione levels (Atkuri et al., 2007; Dean et al., 2011). Spint1a-deficient larvae treated from 1-3 dpf with 100 µM NAC displayed a statistically significant reduction in skin neutrophil infiltration together with reduced skin alterations (Figures 3A, 3B, S2A, S2B). We next tested mito-Tempo, an antioxidant that specifically accumulates in the mitochondria imitating superoxide dismutase activity against superoxide and alkyl radical (Ni et al., 2016), and tempol, a nitroxide antioxidant that acts against the peroxynitrite decomposition compounds, nitrogen dioxide and superoxide radical anion (Mustafa et al., 2015). Whereas both antioxidants were able to reduce skin neutrophil infiltration (Figures 3C, 3D), NFKB activity (Figures 3E, 3F) and morphological alterations (Figures S2C, S2D) in Spint1a-deficient larvae, tempol was found to be much more potent than mito-Tempo (100 µM vs. 100 nM). The relevance of cytosolic rather than mitochondrial ROS was further confirmed by the ability of pharmacological inhibition of NAPDH oxidases with apocynin to alleviate skin neutrophil infiltration of Spint1a-deficient larvae (Figures 3G, 3H) and skin alterations (Figures S2E, S2F). In addition, genetic inhibition of Nox1, Nox4 and Nox5 showed that all contributed to ROS production and skin inflammation (Figures 3I-3M, S2G-S2J, S3). Collectively, these results demonstrate that NADPH oxidase-derived ROS promote skin inflammation in Spint1a-deficient larvae.

**Figure 3.**
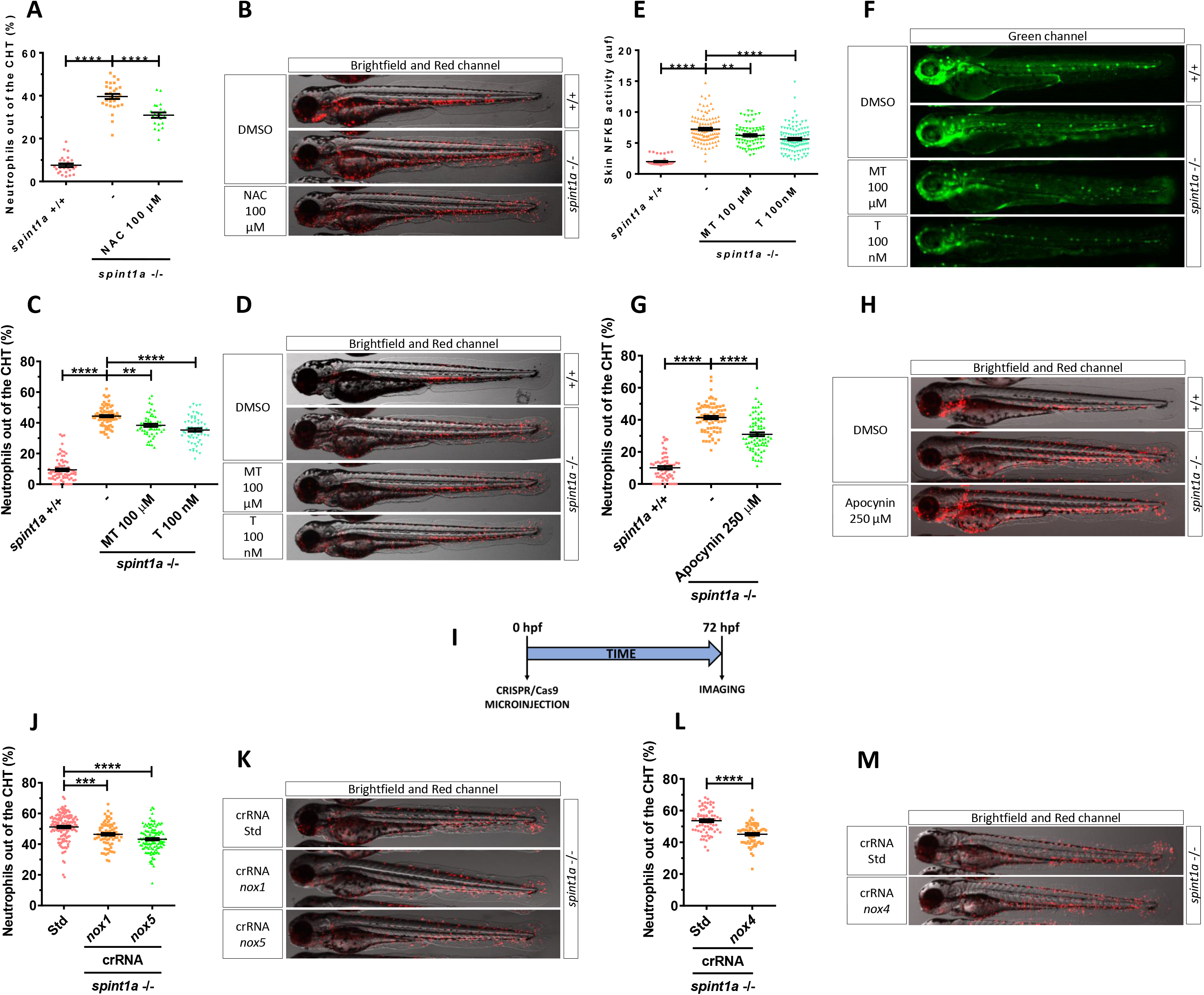
NADPH oxidase-derived ROS promote skin inflammation in Spint1a-deficient larvae. Quantification of neutrophil dispersion out of CHT of wild type and Spint1a-deficient embryos treated with 100 µM N-acetylcysteine (NAC) (A), 100 µM mito-TEMPO (MT) and 100 nM tempol (C), 250 µM apocynin (G), or upon genetic inactivation of *nox1* and *nox5* (I, J) and nox4 (I, M) with CRISPR/Cas9. Representative merge images (brightfield and red channels) of *lyz:dsRED* zebrafish larvae of every group are shown (B, D, F, H, K, M). Determination of NFKB transcriptional activity in the skin of embryos treated with MT and T (E) and representative images (green channel) of *nfkb:eGFP* zebrafish larvae of every group (F). Each dot represents one individual and the mean ± S.E.M. for each group is also shown. P values were calculated using one-way ANOVA and Tukey multiple range test. ns, not significant, **p≤0.01, ****p≤0.0001.

### Inhibition of Parp1 alleviates oxidative stress and skin inflammation of Spint1a-deficient larvae

Due to the pleiotropic roles of NAD^+^ that participates in more than 500 enzymatic reactions and regulates several key cellular processes (Rajman et al., 2018), we analyzed major NAD^+^ consuming enzymes, namely Cd38 and sirtuins (Canto et al., 2015). Pharmacological inhibition of either Cd38 with 78C (Haffner et al., 2015) or sirtuin 1 with EX527 (selisistat) (Napper et al., 2005) were unable to rescue skin inflammation of Spint1a-deficient larvae (data not shown). We then hypothesized that PARPs, which are other NAD^+^-consuming enzymes, could be involved in skin inflammation. Pharmacological inhibition of Parps with olaparib efficiently rescued skin neutrophil infiltration (Figures 4A, 4B), NFKB activation (Figures 4C, 4D) and morphological alterations (Figures S4A, S4B) in Spint1a-deficient larvae. Consistently, genetic inhibition of Parp1 (Figures S4C-S4F) and treatment of larvae with other Parp inhibitors (veliparib and talazoparib) (Figures 4E, 4F) gave similar results, although the inhibitors showed different potencies (500 µM veliparib > 100 µM olaparib >1 µM talazoparib). Furthermore, increased PAR levels were found by western blot in the tail tissue of Spint1a-deficient larvae compared with their wild type siblings (Figure S4G), which could be attenuated by both FK-866 and olaparib (Figure S4H). Similarly, Nampt (FK-866) and Parp (olaparib) inhibition significantly reduced skin epithelial lesions in the *psoriasis* mutant (Figures S5A-S5C), which share the skin inflammation and keratinocyte hyperproliferation phenotypes with the Spint1a-deficient line but has a loss-of-function mutation in *atp1b1a*, which encodes the beta subunit of a Na,K-ATPase pump (Hatzold et al., 2016; Webb et al., 2008).

**Figure 4.**
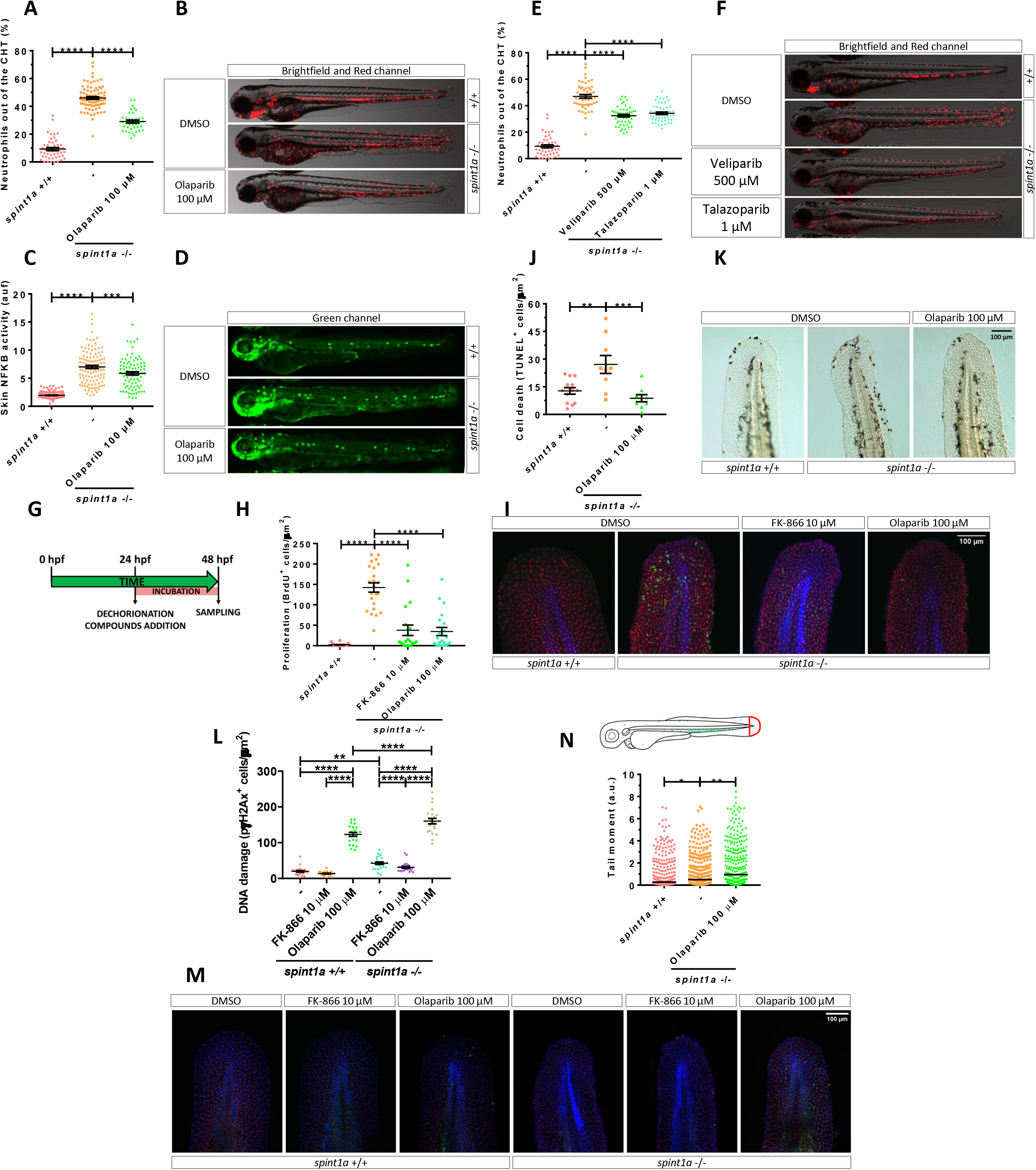
Pharmacological inhibition of Nampt and Parp1 alleviates skin oxidative stress and inflammation, and keratinocyte cell death, hyperproliferation and DNA damage in Spint1a-deficient larvae. Analysis of neutrophil distribution (A, E) and NFKB transcriptional activity in the skin (C) of wild type and Spint1a-deficient larvae larvae treated with olaparib (A, C), and veliparib or talazoparib (E). Representative images (brightfield and red channel in B, F; green channel in D) of *lyz:dsRED* and *nfkb:eGFP* zebrafish larvae of every group are shown. Determination of BrdU positive cells from 48 hpf wild type and Spint1a-deficient zebrafish larvae treated for 24 hours with 10 µM FK-866 or 100 µM olaparib (G, H). Representative merge images maximum intensity projection of a confocal Z stack from zebrafish larvae of every group are shown (I). WIHC with anti-BrdU (green), anti-p63 (red, basal keratinocyte marker) were counterstained with DAPI (blue). Quantification of TUNEL positive cells from 48 hpf wild type and Spint1a-deficient zebrafish larvae treated for 24 hours with 100 µM Olaparib (J). Representative images of zebrafish larvae of every group are shown (K). Quantification of pγH2Ax positive cells from 48 hpf wild type and Spint1a-mutant zebrafish larvae treated for 24 hours with 10 µM FK-866 or 100 µM olaparib (L). Similarly, around 60 zebrafish tail folds (red boxed area) were amputated and disaggregated into cells for comet assay analysis in alkaline conditions (N). Representative merge images of maximum intensity projection of an apotome Z stack from zebrafish larvae of every group are shown (M). WIHC with anti-pγH2Ax (green), anti-P63 (basal keratinocyte marker, red) were counterstained with DAPI (blue). Scale bars, 100 µm. Each dot represents one individual. The mean ± S.E.M. (A-L) and median (N) for each group is shown. P values were calculated using one-way ANOVA and Tukey multiple range test (A-L) and Kruskal-Wallis test and Dunn’s multiple comparisons test (N). *p≤0.05, **p≤0.01, ***p≤0.001, ****p≤0.0001.

### Inhibition of Nampt and Parp1 dampens keratinocyte hyperproliferation and cell death of Spint1a-deficient larvae

As the Spint1a-deficient phenotype starts with basal keratinocyte aggregation, mesenchymal-like properties acquisition and cell death, leading to uncontrolled proliferation (Carney et al., 2007; Mathias et al., 2007), we next analyzed whether Nampt or Parp1 inhibition affected keratinocyte proliferation and/or cell death. As early as 24 hours of treatment, pharmacological inhibition of either Nampt or Parps robustly reduced keratinocyte hyperproliferation in Spint1a-deficient larvae, assayed as BrdU incorporation (Figures 4G-4I), confirming the ability of both Nampt and Parp inhibition to reduce keratinocyte aggregates. Unexpectedly, olaparib also reduced the high number of terminal deoxynucleotidyl transferase dUTP nick end labeling (TUNEL)^+^ keratinocytes found in Spint1a-deficient larvae (Figures 4J, 4K), despite the fact that Parp1 inhibition is expected to lead to the accumulation of DNA lesions and eventually to cell death (Schreiber et al., 2006). We, therefore, investigated DNA damage analyzing the presence of the phosphorylated histone variant H2AX (pγH2Ax), which label dsBs and is independent of PARP1 (Schreiber et al., 2006), and performing a comet assay, which under alkaline conditions can detect both ssBs and dsBs (Pu et al., 2015). The results showed higher keratinocyte DNA damage, assayed by either pγH2Ax staining (Figures 4L, 4M) or by comet assay (Figure 4N), in Spint1a-deficient keratinocytes. In line with previous studies, inhibition of Parp activity with olaparib increased DNA damage in wild type and Spint1a-deficient larvae (Figures 4L, 4M). Remarkably, olaparib-induced DNA lesions were higher in Spint1a-deficient larvae (Figures 4L, 4N), suggesting increased susceptibility to Parp inhibition. Another interesting result was the reduced pγH2Ax staining in Spint1a-deficient keratinocytes in response to FK-866 treatment (Figures 4L, 4M). Taken together, these results suggest that Spint1a-deficient keratinocytes accumulate DNA breaks and have increased susceptibility to DNA stressors.

### Inhibition of parthanatos rescues skin inflammation of Spint1a-deficient larvae

Although the initial characterization of the Spint1a-deficient line revealed high keratinocyte cell death, it was refractory to inhibition of caspase or pro-apoptotic factors, suggesting an unidentified programmed cell death pathway (Carney et al., 2007; Mathias et al., 2007). Similarly, we did not find active caspase-3^+^ keratinocytes in 2 dpf Spint1a-deficient or wild type larvae (Figures 5A-5C). In addition, FK-866 or olaparib treatments did not induce apoptosis (Figures 5A-5C).

**Figure 5.**
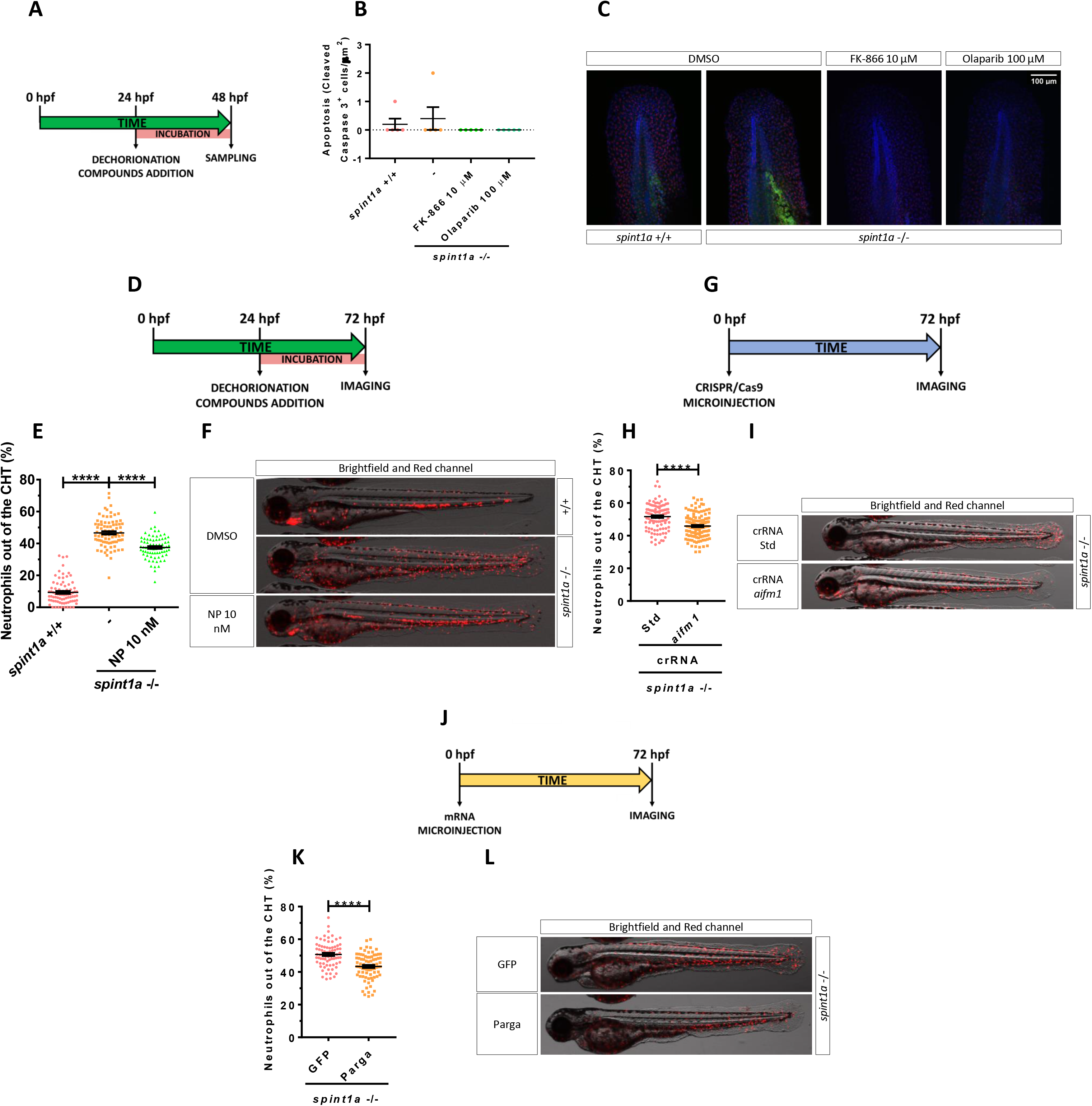
Inhibition of parthanatos rescues skin inflammation of Spint1a-deficient larvae. Quantification of cleaved caspase 3 positive cells from 48 hpf wild type and Spint1a-deficient larvae treated for 24 hours with 10 μM FK-866 or 100 μM Olaparib (A, B). Representative merge images of maximum intensity projection of an apotome Z stack from zebrafish larvae of every group are shown (C). WIHC with anti-cleaved Casp3 (green), anti-P63 (basal keratinocyte marker, red) were counterstained with DAPI (blue) (C). Pharmacological and genetic inhibition experimental settings (D, G). Quantification of the percentage of neutrophils out of the CHT in embryos treated with 10 nM NP (Aifm1 translocation inhibitor) (E), *aifm1* genetic inhibition (H) and *parga* mRNA overexpression (K). Representative merge images (brightfield and red channels) of *lyz:dsRED* zebrafish larvae of every group are shown (F, I, L). For mRNA overexpression, one-cell stage zebrafish eggs were microinjected and imaging was performed in 3dpf larvae (J). Each dot represents one individual and the mean ± S.E.M. for each group is also shown. P values were calculated using one-way ANOVA and Tukey multiple range test. ns, not significant, ****p≤0.0001.

The absence of apoptosis in Spint1a-deficient larvae, together with the ability of Parp inhibitors to robustly reduce keratinocyte cell death in this model, led us to hypothesize that Parp1 overactivation in response to extensive ROS-mediated DNA damage would promote parthanatos cell death. To test this hypothesis, we used N-phenylmaleimide (NP), a chemical inhibitor of Aifm1 translocation from the nucleus to the mitochondria (Susin et al., 1996). Treatment of Spint1a-deficient larvae with 10 nM NP showed statistically significant reduced skin neutrophil infiltration (Figures 5D-5F) and keratinocyte aggregates (Figures S6A, S6B). Similar results were found upon genetic inhibition of Aifm (Figures 5G-5I and S6C, S6D) and forced expression of *parga*, which encodes PAR glycohydrolase a (Figures 5J-5L and S6E-S6G). Collectively, these results further confirm that overactivation of Parp1 in Spint1a-deficient animals mediates keratinocyte cell death through parthanatos rather than via depletion of ATP and NAD^+^ cellular stores.

### Pharmacological inhibition of NADPH oxidases/NAMPT/PARP/AIFM1 axis decreased expression of pathology-associated genes in human organotypic 3D skin models of psoriasis

The results obtained in zebrafish prompted us to study the impact of inhibition of oxidative stress, NAMPT and PARP1 in organotypic 3D human psoriasis models (Figure 6A). The results showed robust increase of NAMPT transcript levels in psoriatic epidermis, i.e. stimulated with cytokines IL17 and IL22 (Figure 6B). Pharmacological inhibition of NADPH oxidases with apocynin significantly reduced the mRNA levels of the inflammation marker defensin β4 (*DEFB4*) (Figure 6A), while it did not affect those of the differentiation markers filaggrin (*FLG*) and loricrin (*LOR*) (Figure 6C). Similarly, inhibition of NAMPT, PARP and AIFM1 reduced the transcript levels *DEFB4* and *S100A8*, another inflammation marker-associated to psoriasis (Figure 6B), while all treatments further reduced LOR and FLR mRNA levels, as did all-trans retinoic acid (ATRA) (Figure 6C). These results confirmed that inhibition of parthanatos cells death with NADPH oxidases, NAMPT, PARP and AIFM1 inhibitors also reduced inflammation in human psoriasis models.

**Figure 6.**
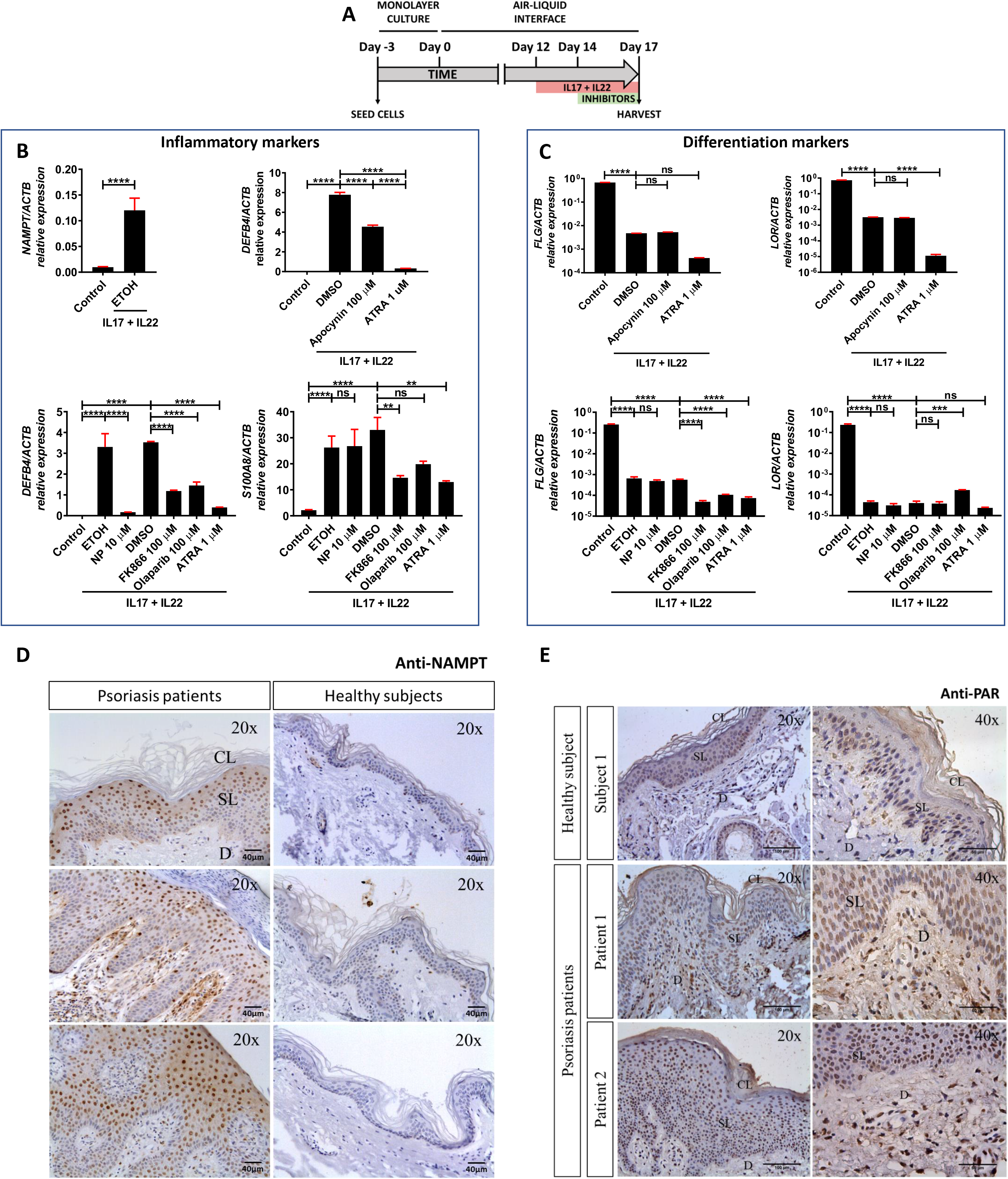
Reduction of pathology-associated genes expression in human psoriatic epidermis organoids by inhibition of NADPH oxidases/NAMPT/PARP/AIFM1 axis, and robust increase of NAMPT protein amount and PARylation in lesional skin from psoriasis patients. (A-C) Transcript levels of the indicated genes encoding inflammation (B) and differentiation (C) markers were determined in human organotypic 3D skin models pre-treated with 30 ng/ml IL17A and IL22 in the presence of vehicles (ETOH and DMSO) or the indicated inhibitors. The mean ± S.E.M. for each group is shown. The results are representative of 3 independent experiments. P values were calculated using one-way ANOVA and Tukey multiple range test. ****p≤0.0001, *****p ≤0.00001. (C, D). Representative images of sections from healthy and psoriatic skin biopsies that have been immunostained with an anti-NAMPT monoclonal antibody (E) or anti-poly (ADP-ribose) monoclonal antibody (F) and then slightly counterstained with hematoxilin. Scale bar is 40 µm in E and 100 µm and 50 µm in left and right panels, respectively, in F. CL, Cornified layer; SL, Spinous layer; D, dermis.

### The expression of genes encoding NAD^+^ and PAR metabolic enzymes is altered in psoriasis in humans

Transcriptomic analysis of human psoriasis lesional skin data collected in the GEO database revealed differential expression profile of genes encoding enzymes involved in NAD^+^ and PAR (Figures S7, S8) metabolism. The mRNA levels *NAMPT* were upregulated (Figure S8), while those of *NMNAT3* were downregulated in psoriasis lesional skin (Figures S8). The transcript levels of *NRK2*, which is involved NR conversion (Nikiforov et al., 2015), were upregulated in psoriasis lesional skin (Figures S8). Preiss-Handler pathway seemed to show an intensified activity only in psoriasis lesional skin, since upregulation of the mRNA levels of genes encoding its components *NAPRT*, encoding nicotinate phosphoribosyltransferase, and *NADSYN*, encoding NAD synthetase, was observed (Figures S8). Finally, the genes coding for NAD^+^ biosynthetic enzymes involved in *de novo* pathway were also altered: while the transcript levels of *IDO1*, encoding indoleamine 2,3-dioxygenase 1, and *TDO2*, encoding tryptophan 2,3-dioxygenase, were induced in lesional skin, *QPRT*, encoding quinolinate phosphoribosyltransferase, slightly decreased in psoriasis compared to healthy skin (Figures S7A). As regards genes encoding enzymes involved in NAD^+^ degradation, increased *CD38* transcript levels were observed in psoriasis lesional skin (Figures S8).

We next analyzed the transcript levels of *PARP1*, *AIFM1* and *MIF*, which encode three indispensable parthanatos components (Wang et al., 2016) as well as those encoding different PAR hydrolases that negatively regulate protein PARylation. Transcriptomic data revealed strong increased mRNA levels of *PARP1*, *AIFM1* and *MIF* in psoriasis lesional skin (Figures S9). Although no alteration in the expression profile of the gene encoding *PARG* was found, the transcript levels of the genes encoding several PAR hydrolases, namely *MACROD1*, *MACROD2* and *TARG1* were lower in psoriasis lesional skin (Figures S9). Furthermore, genes encoding other PAR hydrolases were specifically upregulated in psoriasis lesional skin (*ARH3*) or downregulated (*NUDT16* and *ENPP1*) (Figures S9). In addition, psoriasis lesional skin also showed enhanced transcript levels of *ARH1* (Figures S9), whose product cleaves the terminal bond but only for targets PARylated on arginine (Qi et al., 2019). These results taken together indicate that psoriasis may display increased PARylation.

### NAMPT and PAR increase in the nucleus of human keratinocytes from psoriasis lesions

Immunohistochemical analysis of samples from healthy skin and psoriasis lesions showed that NAMPT was hardly detected in healthy epidermis and dermis (Figure 6D). However, NAMPT was widely overexpressed in the spinous layer and in a few basal keratinocytes and dermal cells in psoriasis lesional skin (Figure 6D). Curiously, NAMPT immunoreactivity was mainly found in the nucleus of keratinocytes but a fainter immunoreactivity was also observed in their cytoplasm (Figure 6D). Consistently, although PAR immunoreactivity was found in the nuclei of scattered keratinocytes of the spinous layer from healthy skin subjects, it was widely observed in the nuclei of most keratinocytes of the spinous layer and dermal fibroblasts from psoriasis lesional skin (Figure 6E), demonstrating increased PARylation in psoriasis lesional skin.

## Discussion

NAD^+^ metabolism plays a fundamental role in maintaining organism homeostasis. NAMPT, the rate-limiting step enzyme in the NAD^+^ salvage pathway, has been associated to oxidative stress and inflammation (Garten et al., 2015), being identified as a universal biomarker of chronic inflammation, including psoriasis (Mesko et al., 2010). PARPs are major NAD^+^ consuming enzymes involved in DNA repair, although their involvement in inflammation has also been widely recognized (Canto et al., 2015; Qi et al., 2019). We have shown here using the unique advantages of the zebrafish embryo/larval model for *in vivo* imaging and drug/genetic screening a crucial contribution of NAMPT and PARP1 to skin inflammation through the induction of parthanatos cell death (Figure 7). Consistently, inhibition of parthanatos also alleviated inflammation in human organotypic 3D models of psoriasis. To the best of our knowledge, this is the first study demonstrating the existence of parthanatos *in vivo*. In these models, the ability of NAD^+^ and its precursors to induce oxidative stress can be explained by their capacity to boost NADH/NADPH intracellular levels that would fuel NADPH oxidases to generate ROS (Figure 7). Consistently, pharmacological and genetic inhibition of NADPH oxidases or NAMPT efficiently counteracted skin H2O2 synthesis and inflammation in both zebrafish and human models. The relevance of oxidative stress in psoriasis has not been demonstrated definitively but psoriasis patients show increased levels of oxidative stress markers, decreased levels of antioxidant molecules and reduced activity of main antioxidant enzymes, such as superoxidase dismutase and catalase (Lin and Huang, 2016; Nemati et al., 2014). Additionally, high levels of oxidized guanine species, a marker of DNA/RNA damage, were found in the serum of psoriasis patients (Borska et al., 2017), further highlighting the relevance of the Spint1a-deficient line to study psoriasis. Curiously, high doses of FK-866 triggered NFKB and neutrophil infiltration into the muscle, probably reflecting disruption of the cellular bioenergetics in this tissue due to critically low NAD^+^ levels. This would need further investigation but it is out of the scope of this study.

**Figure 7.**
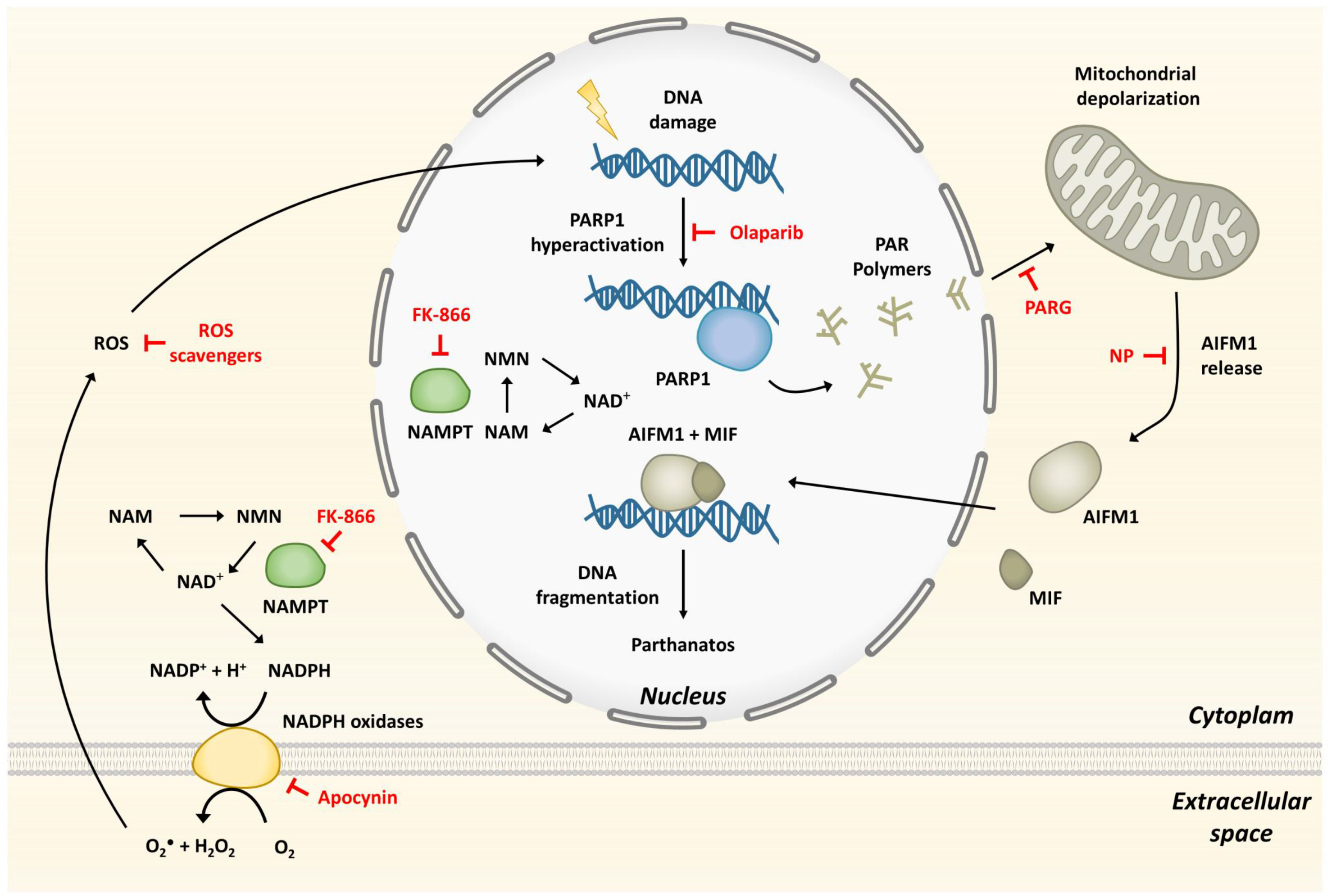
Model showing that hyperactivation of Parp1 in response to ROS-induced DNA damage, and fuelled by NAMPT-derived NAD+, mediates inflammation through parthanatos. The inhibitors used are depicted in red.

We also revealed new features of the Spint1a-deficient zebrafish line that may be useful in future studies. This model is of relevance for the study of human skin inflammatory disorders, since exacerbated serine protease activity is responsible for skin barrier defects and chronic inflammation in Netherton syndrome, a rare but severe autosomal recessive form of ichthyosis caused by mutation in *SPINK5*, which encodes the serine protease inhibitor LEKTI (Sarri et al., 2017). Notably, ablation of matriptase from LEKTI-deficient mice rescues skin barrier defects and inflammation (Sales et al., 2010). Furthermore, mutations in *SPINK5* have also been associated with atopic dermatitis (Liang et al., 2016), further highlighting the relevance of the Spint1a-deficient zebrafish model. We found that this model exhibited increased H2O2 release in the skin, a well-known signal for leukocyte recruitment to acute (Niethammer et al., 2009) and chronic (Candel et al., 2014) insults. H2O2 was released at high levels in keratinocyte aggregates; precisely where higher NFKB levels were also observed and neutrophils were actively recruited. The relevance of oxidative stress in the skin inflammation of this line was confirmed by ROS scavenging and inhibition of NADPH oxidases and Nampt, being all able to restore skin morphology (Figure 7). We also found that Spint1a-deficient larvae exhibited extensive DNA damage, evaluated by pγH2Ax staining and comet assay, and had increased susceptibility to olaparib, a DNA damage inducer (Kukolj et al., 2017). This is an important observation from a clinical point of view, since psoriasis patients receiving PUVA have a significantly increased risk for the development of skin cancers (primarily squamous cell carcinomas) (Gasparro, 2000). Pharmacological inhibition of Nampt not only reduced oxidative stress, skin inflammation and keratinocyte hyperproliferation but also DNA damage. However, inhibition of Parp1 with olaparib increased keratinocyte DNA damage, despite being able to reduce at the same time cell death and PARylation. Moreover, neither olaparib nor FK-866 treatments promoted keratinocyte apoptosis, contrasting previous studies where PARP inhibitors were reported to induce apoptosis in proliferating cells (Schreiber et al., 2006). Therefore, olaparib induced an undesirable situation from a clinical point of view, where keratinocytes accumulated DNA damage but did not suffer apoptosis or another form of cell death. This is not surprising, since psoriatic keratinocytes are highly resistant to apoptosis (Kastelan et al., 2009) and its induction by PUVA may be involved in its therapeutic effects (Laporte et al., 2000).

The depletion of NAD^+^ cellular stores by Nampt inhibition might have also resulted in a direct inhibition of keratinocyte proliferation in the Spint1a-deficient and human organotypic 3D skin models of psoriasis. Indeed, it is known that cancer cells are more sensitive to the loss of NAMPT, a distinctive feature that was proposed to be used clinically in combination with other anticancer agents (Garten et al., 2015). Moreover, various treatments for psoriasis target proliferation of keratinocytes and immune cells, such as acitretin, PUVA, methotrexate and cyclosporine, among others (Greaves and Weinstein, 1995). However, no developmental alterations or delay were observed in larvae treated with Nampt or Parp inhibitors in 3 dpf larvae. Therefore, although these inhibitors caused a blockage in cell proliferation, the results are consistent with a recovery of the mutant phenotype by the inhibition of oxidative stress and parthanatos.

One of the most interesting observations of this study is that Parp1 inhibition was able to restore epithelial homeostasis and to alleviate skin inflammation in both Spint1a- and Atp1b1a-deficient models. The ability of Aifm1 inhibition and Parga overexpression to phenocopy the effects of Nampt and Parp inhibition in the Spint1a-deficient model further confirmed that parthanatos induced by Parp1 overactivation, rather than by depletion of ATP and NAD^+^ keratinocyte stores, mediated skin inflammation (Figure 7). This is consistent with the results obtained with Nampt inhibition, which resulted in depleted NAD^+^ levels, but was also able to rescue skin inflammation and keratinocyte cell death and hyperproliferation in both zebrafish models. However, Nampt inhibition would simultaneously hamper NADPH oxidases and Parp1 enzymatic activities, while Parp inhibition would only suppress the latter. Anyway, prevention of cell death by either strategy might reduce danger signal and cytokine release, limiting immune cell recruitment and dampen inflammation (Kunze and Hottiger, 2019) by blocking the inflammatory loop that propagates in psoriasis (Dainichi et al., 2018).

The analysis of human transcriptomic data from psoriasis lesional skin showed altered expression profiles of genes encoding NAD^+^ metabolic enzymes, including salvage, Preiss-Handler and de novo pathways, and of genes encoding key enzymes involved in PAR metabolism and parthanatos. As both diseases are characterized by an important cellular immune infiltration and growth of other cell types like nerves or blood vessels, the genes whose expression is altered in lesional skin may be expressed in different tissues other than keratinocytes. Our immunohistochemical analysis of psoriasis lesional skin confirmed the drastic induction of NAMPT at protein level and PAR accumulation in the nucleus of epidermal keratinocytes and dermal cells.

In conclusion, we report that hyperactivation of Parp1 in response to ROS-induced DNA damage, and fuelled by NAMPT-derived NAD^+^, mediates inflammation through parthanatos cell death in preclinical zebrafish and human organotypic 3D skin models of psoriasis. The altered expression of genes encoding key enzymes involved in NAD^+^ and PAR metabolism in psoriasis lesional skin, and in particular the robust induction of NAMPT and the accumulation of PAR in psoriasis lesional skin, point to NAMPT, PARP1 and AIFM1 as novel therapeutic targets to treat psoriasis and probably other skin inflammatory disorders.

## Methods

### Ethics statement

The experiments performed comply with the Guidelines of the European Union Council (Directive 2010/63/EU) and the Spanish RD 53/2013. Experiments and procedures were performed as approved by the Bioethical Committees of the University of Murcia (approval numbers #75/2014, #216/2014 and 395/2017) and Ethical Clinical Research Committee of The University Hospital Virgen de la Arrixaca (approval number #8/13).

### Animals

Wild-type zebrafish (*Danio rerio* H.) lines AB, TL and WIK obtained from the Zebrafish International Resource Center (ZIRC) were used and handled according to the zebrafish handbook (Westerfield, 2000). The transgenic zebrafish line *Tg(lyz:dsRED2)^nz50^* (Hall et al., 2007) and Tg(NFκB-RE:eGFP) (Kanther et al., 2011) were described previously. The mutant zebrafish lines *spint1a^hi2217Tg/hi2217Tg^* (Amsterdam et al., 1999) and the *atpb1a^m14/m14^* (Webb et al., 2008) were isolated from insertional and ehyl methanesulfonate induced mutagenesis screens, respectively.

### Genetic inhibition in zebrafish

The crispr RNA (crRNA) obtained from Integrated DNA technologies (IDT) with the following target sequence were used: *aifm1* crRNA: 5’-CTTGCCAAGGTGGAGAACGG-3’, *parp1* crRNA: 5’-TGGATTTACTGACCTCCGCT-3’, *nampta* 5’-AGTAAAGAGCACATTTCCCCG-3’; *namptb* 5’-GGAGTAGACTTTATTTATAT-3’; *nox1* crRNA: CAAGCTGGTGGCCTACATGA; *nox4* crRNA: TTCGCTTGTGTCCTTCAAGC and *nox5* crRNA: GAGGTCATGGAAAATCTCAC. They were resuspended in duplex buffer at 100 µM and 1 µl was incubated with 1 µl of 100 µM trans-activating CRISPR RNA (tracrRNA) at 95°C for 5 minutes and then 5 minutes at room temperature to form the complex. One µl of this complex was mixed with 0.25 µl of recombinant Cas9 (10 mg/ml) and 3.75 µl of duplex buffer. The efficiency of each crRNA was determined by the TIDE webtool (https://tide.deskgen.com/) (Brinkman et al., 2014) or T7 nuclease assay. Crispant larvae of 3 dpf were used in all studies.

### Chemical treatments in zebrafish

Zebrafish embryos were manually dechorionated at 24 hpf. Larvae were treated from 24 hpf to 48 hpf or 72 hpf by chemical bath immersion at 28 °C. Incubation was carried out in 6-or 24-well plates containing 20-25 larvae/well in egg water (including 60 µg/mL sea salts in distilled water) supplemented with 1% dimethyl sulfoxide (DMSO). The inhibitors and metabolites used, the concentrations tested and their targets are shown in Table S1.

### Imaging of zebrafish larvae

Live imaging of 72 hpf larvae was obtained employing buffered tricaine (200 µg/mL) dissolved in egg water. Images were captured with an epifluorescence LEICA MZ16FA stereomicroscope set up with green and red fluorescent filters. All images were acquired with the integrated camera on the stereomicroscope and were analyzed to determine number of neutrophils and their distribution in the larvae. The transcriptional activity of NF-κB was visualized and measured with the zebrafish line NFκB-RE:eGFP. H2O2 release was quantified employing the live cell fluorogenic substrate acetyl-pentafluorobenzene sulphonyl fluorescein (Cayman Chemical) (Candel et al., 2014; de Oliveira et al., 2015). Briefly, about 20 embryos of 72 hpf were rinsed with egg water and collected in a well of a 24-well plate with 50 µM of the substrate in 1% DMSO for 1 hour. ImageJ software was employed to determine mean intensity fluorescence of a common region of interest (ROI) placed in the dorsal fin for H2O2 production quantification. Similarly, a ROI located in muscle or skin was used to obtain mean intensity fluorescence of NFκB-RE:eGFP transgenic line.

### Whole-mount immunohistochemistry in zebrafish

BrdU incorporation assay was used to determine cell proliferation. Embryos of 48 hpf were incubated in 10 mM of BrdU dissolved in egg water for 3 hours at 28 °C followed by a one-hour wash out with egg water and fixation in 4% paraformaldehyde (PFA) overnight at 4°C or 2 hours at room temperature (RT). For the rest of immunofluorescence techniques, embryos/larvae were directly fixed in 4% PFA, as indicated above. Embryos/larvae were then washed with phosphate buffer saline (PBS) with 0.1% tween-20 (PBST) 3 times for 5 minutes. In order to dehydrate the sample progressively, 25%/75% methanol (MeOH)/PBST, 50%/50% MeOH/PBST and 75%/25% MeOH/PBST and 100% MeOH were employed each for 5 minutes. At this point, embryos were stored at −20 °C. To proceed with the immunofluorescence, samples were re-hydrated in decreasing solutions of MeOH/PBST, as previously described, and then washed 3 times for 5 minutes with PBST. For BrdU staining next step consisted on applying 30 minutes a solution of 2N HCl in PBS supplemented with 0.5 % Triton X-100 (PBSTriton), followed by 3 wash with PBSTriton for 15 minutes. Blocking step was carried out for at least 2 hours at RT with a PBSTriton solution supplemented with 10 % fetal calf serum (FBS) and 0.1 % DMSO. The primary antibody incubation in blocking solution was done overnight at 4 °C or 3-4 hours at room temperature. After that, larvae were washed 6 times for 5 minutes. Incubation in secondary antibody in blocking solution was performed for 2-3 hours in the dark. In order to remove unbound secondary antibody, embryos were washed 3 times for 10 minutes with PBST. In this step, the sample was ready for 2-(4-Amidinophenyl)-6-indolecarbamidine (DAPI) staining with a DAPI solution (1:1000) in PBST for 20 minutes followed by a wash out step of 3 times for 10 minutes with PBST. Finally, embryos were transferred to 80% glycerol/20% PBST and stored in the dark at 4 °C until imaging.

The following primary antibodies were used: rabbit anti-BrdU (Abcam, ab152095, 1:200), mouse anti-p63 (Santa Cruz Biotechnology, sc-7255, 1:200), rabbit Anti-Active Caspase-3 (Bd Bioscience, #559565, 1:250) and rabbit anti-H2AX.XS139ph (phospho Ser139) (GeneTex, GTX127342, 1:200). Secondary antibodies were goat anti-rabbit Alexa Fluor-488 (Molecular probes, CAT#A11008, 1:1000) and goat anti-mouse Cyanine 3 (Life technologies, A10521, 1:1000). Images for BrdU staining were taken using a Zeiss Confocal (LSM710 META), the other stains were acquired by ZEISS Apotome.2. All images were processed using ImageJ software.

### TUNEL Assay

Emrbyos/larvae were fixed and dehydrated as described above. Afterwards, embryos/larvae were rinsed with pre-cooled (−20 °C) 100% acetone and then incubated in 100% acetone at −20 °C for 10 minutes. Samples were then washed 3 times for 10 minutes with PBST and incubated in a solution of 0.1% TritonX-100 and 0.1% sodium citrate (10%) in PBS for 15 minutes to further permeabilize the embryos/larvae. Next step consisted of rinsing specimens 2 times for 5 minutes in PBST. Following complete removal of PBST from samples, it was added 50 µL of fresh TUNEL reaction mixture composed of 5 µL of enzyme solution mixed with 45 µL of labeling solution (In Situ Cell Death Detection kit, POD, ROCHE, version 15.0) for 1 hour at 37 °C, followed by 5 wash with PBST for 5 minutes. Blocking step was carried out for at least 1 hour at room temperature with blocking buffer. To proceed with TUNEL assay, blocking buffer was removed and added 50 µL Converter-POD (anti-fluorescein antibody conjugated to peroxidase) for 1 hour at room temperature or overnight at 4 °C on rocker. Embryos were rinsed 4 times for 30 minutes in PBST and incubated in 1 mL of 3,3′-Diaminobenzidine (DAB) solution for 30 minutes in the dark and transferred to a 24 well-plate. Two µL of a fresh 0.3% H2O2 solution was added to initiate peroxidase reaction that was monitored 10-20 minutes followed by rinsing and a wash out step of 2 times for 5 minutes with PBST. Finally, embryos were transferred to 80% glycerol/20% PBST and stored in dark at 4 °C until imaging. Images were acquired by ZEISS Apotome.2 and processed using ImageJ software.

### Comet assay in zebrafish

Zebrafish embryos at 48 hpf were anesthetized in tricaine (200 µg/mL) dissolved in egg water and the end of the fin fold was amputated with a scalpel. Tissues collected from around 60 embryos were pooled, then spun and resuspended in 1 mL PBS. Liberase at 1:65/volume of PBS (Roche, cat # 05401119001) was added and tissues were incubated at 28 °C for 35 minutes, pipetting up and down every 5 minutes. To stop the reaction, FBS was added to a final concentration of 5% in PBS. From now on, samples were kept on ice. Disaggregated fin folds were filtered through a 40 µM filter and washed using PBS + 5% FBS. Cell suspension was centrifuged at 650xg for 5 minutes and resuspended in 50 µL of PBS + 5% FBS. In order to determine cell number, Trypan Blue-treated cell suspension was applied to Neubauer chamber and cell were counted in an inverted microscope. Around 15,000 cells were employed to perform the Alkaline Comet Assay according to the manufacturer’s protocol (Trevigen). Briefly, cells were added in low melting point agarose at 37 °C at a ratio of 1:10 (v/v) and then were placed onto microscope slides. After adhesion at 4 °C for 30 minutes in the dark, slides were immersed in lysis buffer (precooled at 4 °C) overnight at 4 °C. Next, DNA was unwound in alkaline electrophoresis solution pH>13 (200 mM NaOH, 1 mM EDTA) at room temperature for 20 min in dark, followed by electrophoresis run in the same buffer at 25 V (adjusting the current to 300 mA) for 30 minutes. Slides were washed twice in distilled water for 5 minutes and in 70% ethanol for 5 minutes, then they were dried at 37 °C for 30 minutes. Finally, DNA was stained with SYBR™ Green I Nucleic Acid Gel Stain 10,000X (Invitrogen) and images were taken using a Nikon Eclipse TS2 microscope with 10x objective lens. Quantitative analysis of the tail moment (product of the tail length and percent tail DNA) was obtained using CASPLAB software. More than 100 randomly selected cells were quantified per sample. Values were represented as the median of the tail moment of treated cells relative to the median of the tail moment of untreated cells.

### Western blot

Zebrafish embryos at 72 hpf were anesthetized in tricaine (200 µg/mL) dissolved in egg water and the end of the fin fold was amputated with a scalpel. Tissues collected from around 120 embryos were pooled, then spun and resuspended in 80 µL of 10 mM Tris pH 7.4 + 1% Sodium Dodecyl Sulfate (SDS). Samples were then incubated at 95 oC for 5 min with 1400 rpm agitation, followed by maximum speed centrifugation for 5 min. Supernatants were frozen at – 20 °C until proceeding. BCA kit was employed to quantify protein using BSA as a standard. Fin lysates (10 μg) in SDS sample buffer were subjected to electrophoresis on a polyacrylamide gel and transferred to PVDF membranes. The membranes were incubated for 1 h 30 min with TTBS containing 5% (w/v) skimmed dry milk powder and immunoblotted in the same buffer 16 h at 4 °C with the mouse monoclonal antibody to human poly(ADP-ribose) (1/400, ALX-804-220, Enzo). The blot was then washed with TTBS and incubated for 1 h at room temperature with secondary HRP-conjugated antibody diluted 2500-fold in 5% (w/v) skimmed milk in TTBS. After repeated washes, the signal was detected with the enhanced chemiluminescence reagent and ChemiDoc XRS Biorad.

### Total NAD^+^ and NADH determination

Zebrafish embryos at 72 hpf were anesthetized in tricaine (200 µg/mL) dissolved in cold PBS in order to amputate the tail at the end of the yolk sac extension with a scalpel. Tissues from around 120 embryos were pooled and collected in lysis buffer provided by the kit (ab186032, Abcam) according to the manufacturer’s protocol. Tissues were homogenized and centrifuged at 1400 rpm for 5 minutes at 4 °C. Supernatants were collected and centrifuged at maximum speed for 10 minutes at 4 °C. Supernatants were employed to protein quantification with BCA kit using BSA as a standard. To proceed with Total NAD and NADH determination, 50 µg of protein were employed.

### Gene Expression Omnibus (GEO) database

Human psoriasis (accession number: GSD4602) transcriptomic data collected in the GEO database (https://www.ncbi.nlm.nih.gov/geo/). Gene expression plots were obtained using GraphPad Prism Software.

### Immunohistochemistry in human skin samples

Skin biopsies from healthy donors (n = 5) and psoriasis patients (n = 6) were fixed in 4% PFA, embedded in Paraplast Plus, and sectioned at a thickness of 5 µm. After being dewaxed and rehydrated, the sections were incubated in 10 mM citrate buffer (pH 6) at 95 °C for 30 min and then at room temperature for 20 min to retrieve the antigen. Afterwards, steps to block endogenous peroxidase activity and nonspecific binding were performed. Then, sections were immunostained with a 1 1/100 dilution of mouse monoclonal antibodies to NAMPT (sc-166946, Santa Cruz Biotechnology) a poly (ADP-ribose) (ALX-804-220; Enzo Life Sciences) followed by 1/100 dilution of biotinylated secondary antibody followed by ImmunoCruz® goat ABC Staining System (sc-2023, Santa Cruz Biotechnology) according to manufacturer’s recommendations. Finally, after DAB staining solution was added, sections were dehydrated, cleared and mounted in Neo-Mount. No staining was observed when primary antibody was omitted. Sections were finally examined under a Leica microscope equipped with a digital camera Leica DFC 280, and the photographs were processed with Leica QWin Pro software.

### Human organotypic 3D models

Insert transwells (Sigma-Aldrich MCHT12H48) were seeded with 10^5^ human foreskin keratinocytes (Ker-CT, ATCC CRL-4048) on the transwells in 300 µL CnT-PR medium (CellnTec) in a 12 well format. After 48 hours, cultures were switched to CnT-PR-3D medium (CELLnTEC) for 24 hours and then cultured at the air-liquid interface for 17 days. From day 12 to 17 of the air-liquid interphase culture, the Th17 cytokines IL17A (30 ng/mL) and IL22 (30 ng/mL) were added (Smits et al., 2017). Pharmacological treatments were applied from day 14 to 17 and consisted of 100 µM apocynin, 100 µM FK-866, 100 µM olaparib, 10 µM N-Phenylmaleimide and 1 µM ATRA. Culture medium was refreshed every two days. At day 17, the tissues were harvested for gene expression analysis and immunohistochemistry.

### Statistical analysis

Data were analyzed by analysis of variance (ANOVA) and a Tukey multiple range test to determine differences between groups with gaussian data distribution (square-root transformation were employed for percentage data). Non-parametric data were analyzed by Kruskal-Wallis test and Dunn’s multiple comparisons test. The differences between two samples were analyzed by the Student t-test. The contingency graphs were analyzed by the Chi-square (and Fisher’s exact) test.

## Conflict of interest

A patent for the use of parthanatos inhibitors to treat psoriasis and atopic dermatitis has been registered by Universidad de Murcia and Instituto Murciano de Investigación Biosanitaria (#PCT/EP2020/083380).

## Acknowledgments

We warmly thank I. Fuentes and P. Martínez for their excellent technical assistance and the staff of the Dermatology Service of the University Hospital Virgen de la Arrixaca for patient blood collection. We also thank Profs. S.A. Renshaw and P. Crosier for the zebrafish lines. This work was supported by the Spanish Ministry of Science, Innovation and Universities (grants BIO2014-52655-R and BIO2017-84702-R to VM and PI13/0234 to MLC, and PhD fellowship to FJMN), all co-funded with Fondos Europeos de Desarrollo Regional/European Regional Development Funds), Fundación Séneca-Murcia (grants 20793/PI/18 to VM and 19400/PI/14 to MLC) and the University of Murcia (postdoctoral contracts to ABPO and DGM, and PhD fellowship to FJMM). The funders had no role in the study design, data collection and analysis, decision to publish, or preparation of the manuscript.

## Author contributions

VM conceived the study; FJMM, JCS, ABPO, DGM and VM designed the research; FJMM, JCS, FJMN, IC, IMV, JA, JH, ALM, ABPO and DGM performed the research; FJMM, JCS, FJMN, IC, IMV, JA, JH, ALM, TMM, RCV, JL, MH, JCB, AGA, MLC, ABPO, DGM and VM analyzed the data; and FJMM and VM wrote the manuscript with minor contributions from other authors.

---

**Figure S1, related to Figure 2.**
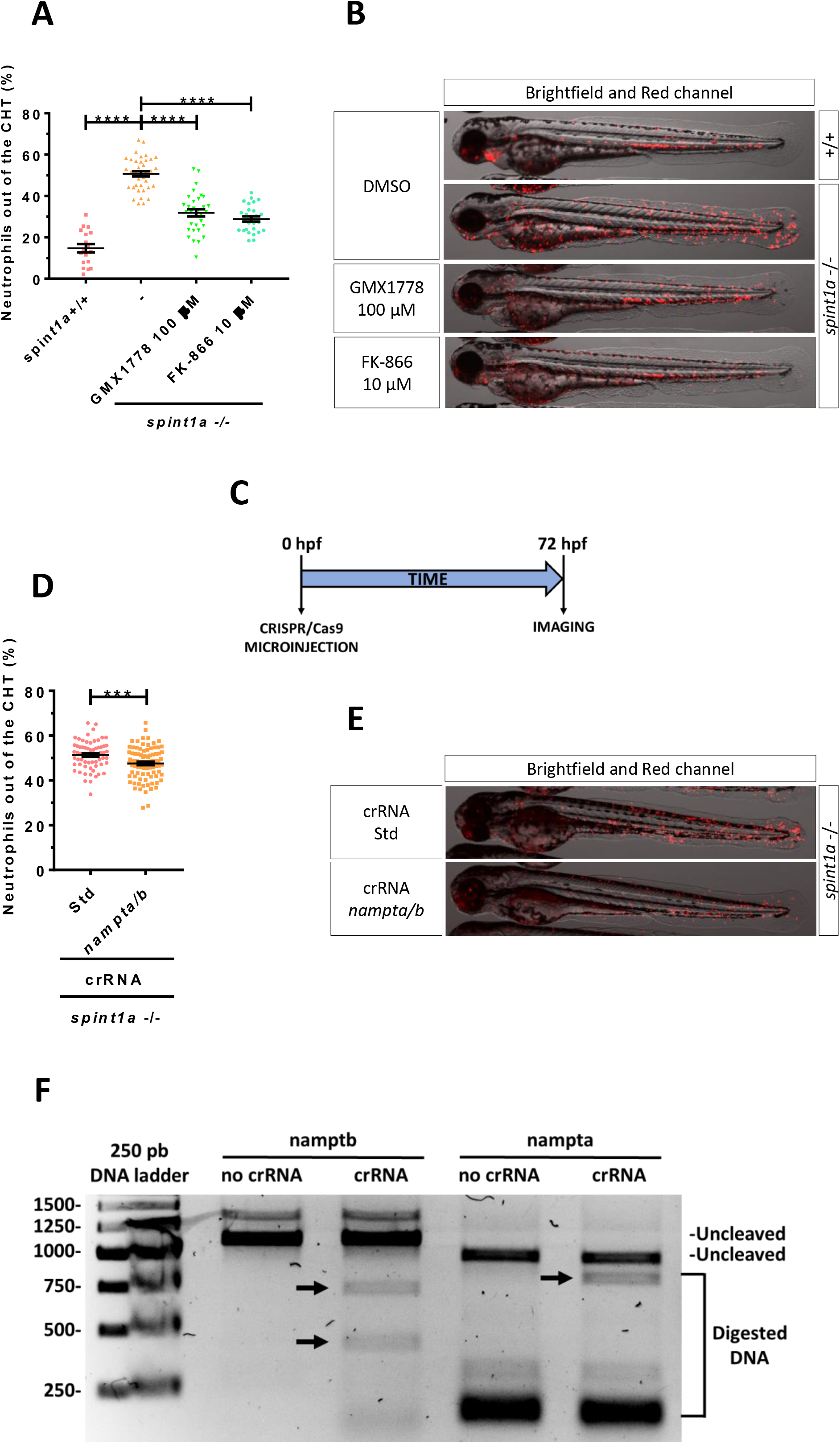
Genetic and pharmacological inhibition of Nampt alleviates skin inflammation and restores epithelial integrity in *spint1a* –deficient larvae. (A) Neutrophil distribution of wild type and Spint1a-deficient larvae treated with the pharmacological inhibitors of Nampt GMX1778 and FK-866. (B) Representative merge images (brightfield and red channels) of *lyz:dsRED* zebrafish larvae of every group are shown. (C) For genetic inhibition using CRISPR/Cas9 technology, one-cell stage zebrafish eggs were microinjected and imaging was performed in 3 dpf larvae (D). Quantification of the percentage of neutrophils out of the CHT in Spint1a-deficient larvae upon knockdown of Nampta/Namptb. (E) Representative merge images (brightfield and red channel) of *lyz:dsRED* zebrafish larvae of every group are shown (F) Agarose gel electrophoresis of *namptb* and *nampta* DNA genomic fragments (arrows) derived from Cas9 *in vitro* digestion in presence or absence of specific *namptb* or *nampta* crRNA. Each dot represents one individual and the mean ± S.E.M. for each group is also shown. P values were calculated using one-way ANOVA and Tukey multiple range test. ***p≤0.001, ****p≤0.0001.

**Figure S2, related to Figure 3.**
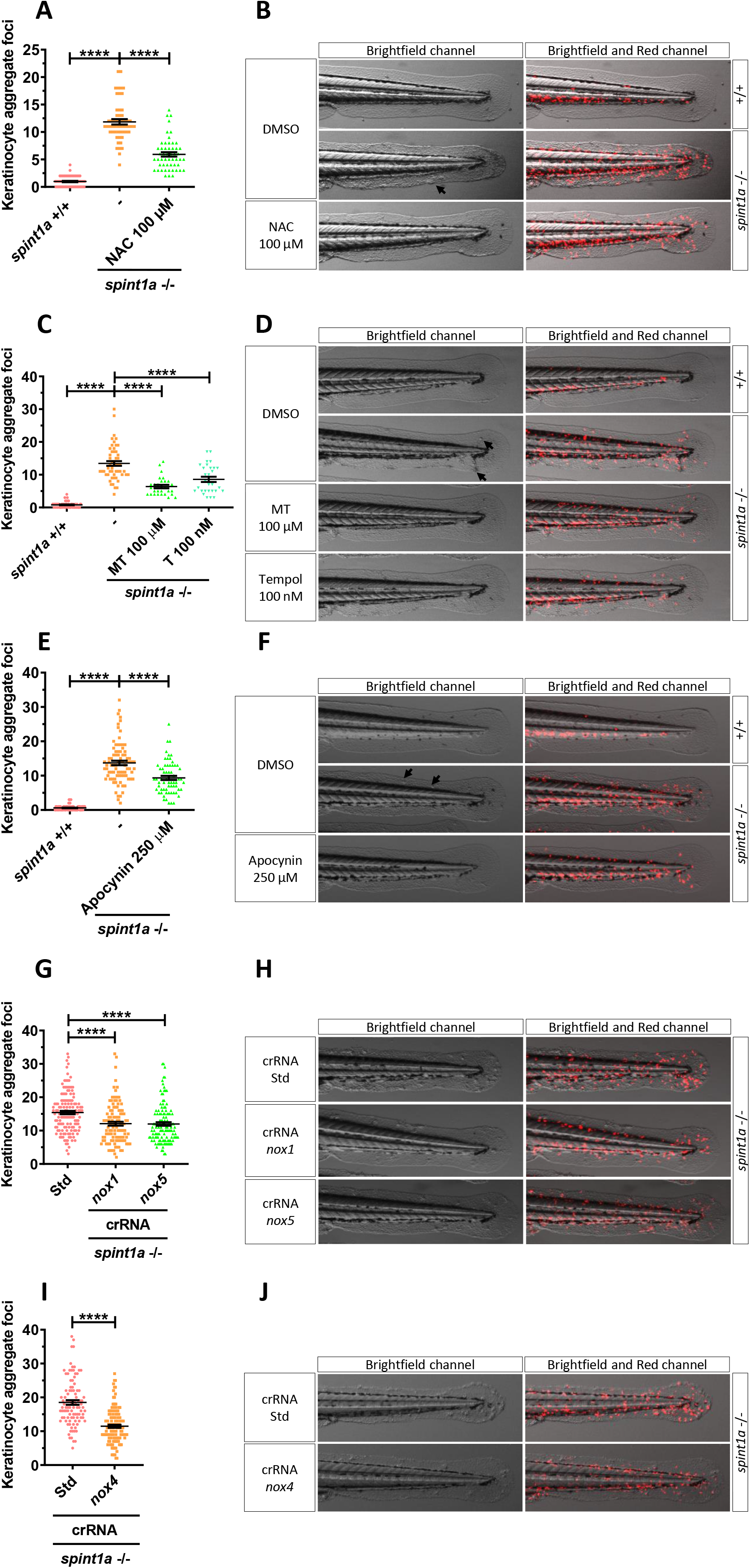
ROS scavenging and inhibition of NADPH-oxidases rescue skin neutrophil recruitment and skin morphological alterations of Spint1-deficient larvae. Quantification of keratinocyte aggregation foci in the tail of *lyz:dsRED* larvae shown in Figure 3 (A, C, E, G, I) and detailed representative merge images (brightfield and red channel) (B, D, F, H, J) upon their treatment with with vehicle (DMSO), 100 µM N-acetylcysteine (NAC) (A, B), 100 µM mito-TEMPO (MT) and 100 nM tempol (T) (C, D), 250 µM apocynin (E, F), or upon genetic inhibition of *nox1* and *nox5* (G, H), and nox4 (I, J).

**Figure S3, related to Figure 3.**
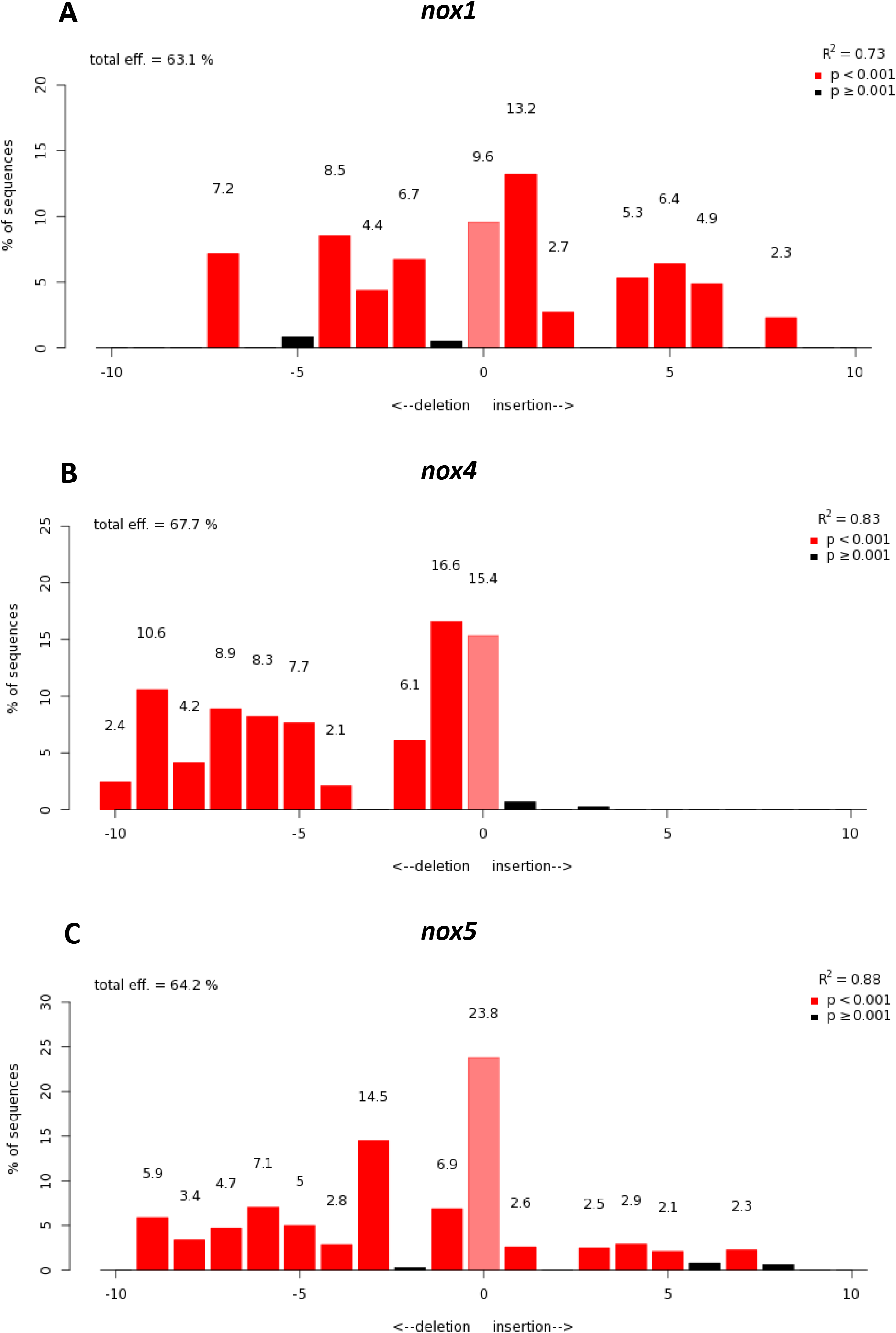
Efficiency of crRNA for *nox1*, *nox4* and *nox5*. Analysis of genome editing efficiency in larvae injected with control or *nox1* (A), *nox4* (B) and *nox5* (C) crRNA/Cas9 complexes and quantification rate of non-homologous end-joining mediated repair (NHEJ) showing all insertions and deletions (INDELs) (https://tide.deskgen.com/).

**Figure S4, related to Figure 4.**
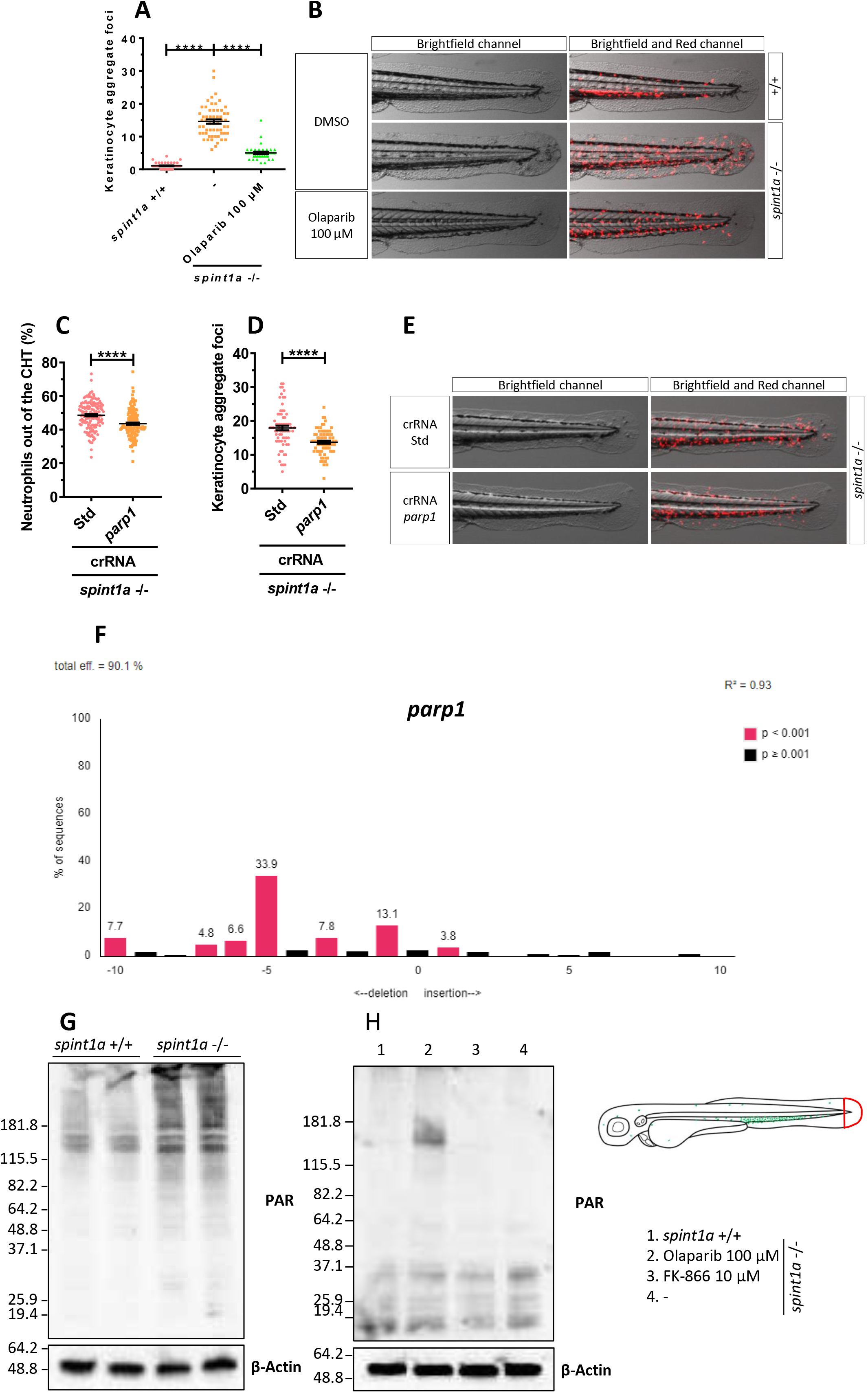
Genetic and pharmacological inhibition of Nampt and Parp1 diminishes PARylation, skin inflammation and restores epithelial integrity in Spint1a-deficient larvae. (A, B) Quantification of keratinocyte aggregates and detailed representative merge images (brightfield and red channels) of wild type and Spint1a-deficient zebrafish treated with vehicle (DMSO) or 100 µM olaparib shown in Figure 4A. (C-E) Neutrophil distribution (C) and keratinocyte aggregates (D) of Spint1a-deficient larvae injected with control or *parp1* crRNA/Cas9 complexes. Representative images are shown in E. (F) Analysis of genome editing efficiency in larvae injected with control or *parp1* crRNA/Cas9 complexes and quantification rate of non-homologous end-joining mediated repair (NHEJ) showing all insertions and deletions (INDELs) (https://tide.deskgen.com/). (G, H) Western blots with anti-PAR and anti-β-actin of tail fold (red boxed area) lysates from 3 dpf wild type and Spint1a-deficient zebrafish larvae treated for 48 hours with 10 µM FK-866 or 100 µM olaparib. Each dot represents one individual and the mean ± S.E.M. for each group is also shown. P values were calculated using one-way ANOVA and Tukey multiple range test. ns, not significant, **p≤0.01, ****p≤0.0001 (C). Representative merge images (brightfield and red channel) of *lyz:dsRED* zebrafish larvae of every group are shown (D).

**Figure S5, related to Figures 2 and 4.**
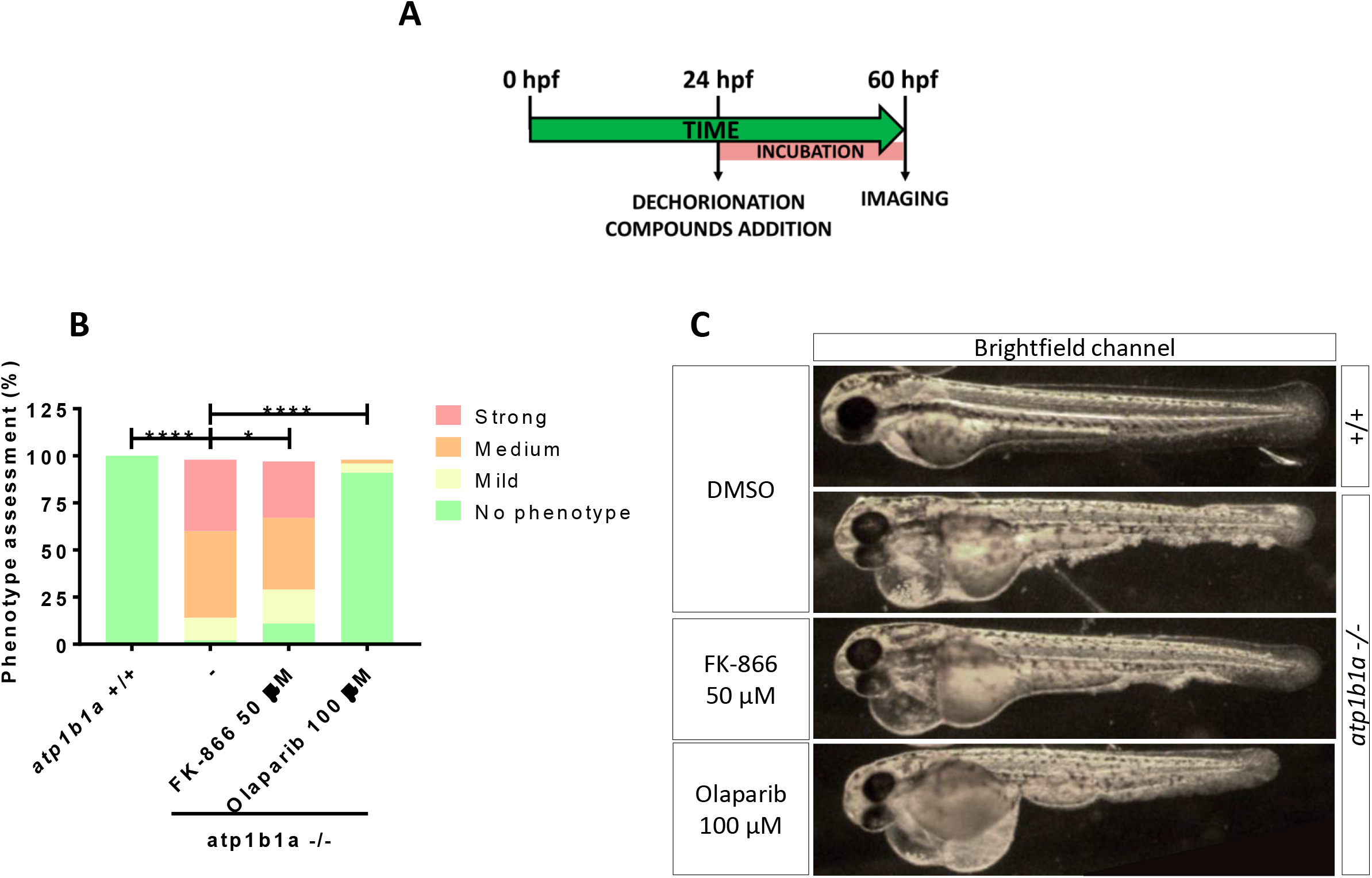
FK-866 and olaparib improve skin epithelial integrity in *psoriasis* mutants. Determination of the skin phenotype of 2.5 dpf zebrafish Atp1b1a-deficient larvae treated 1.5 days with 50 µM FK-866 or 100 µM olaparib (A, B). Representative bright field images of zebrafish larvae of every group are shown (C). P values were calculated using Chi-square and Fisheŕs exact test *p≤0.05, ****p≤0.0001.

**Figure S6, related to Figure 5.**
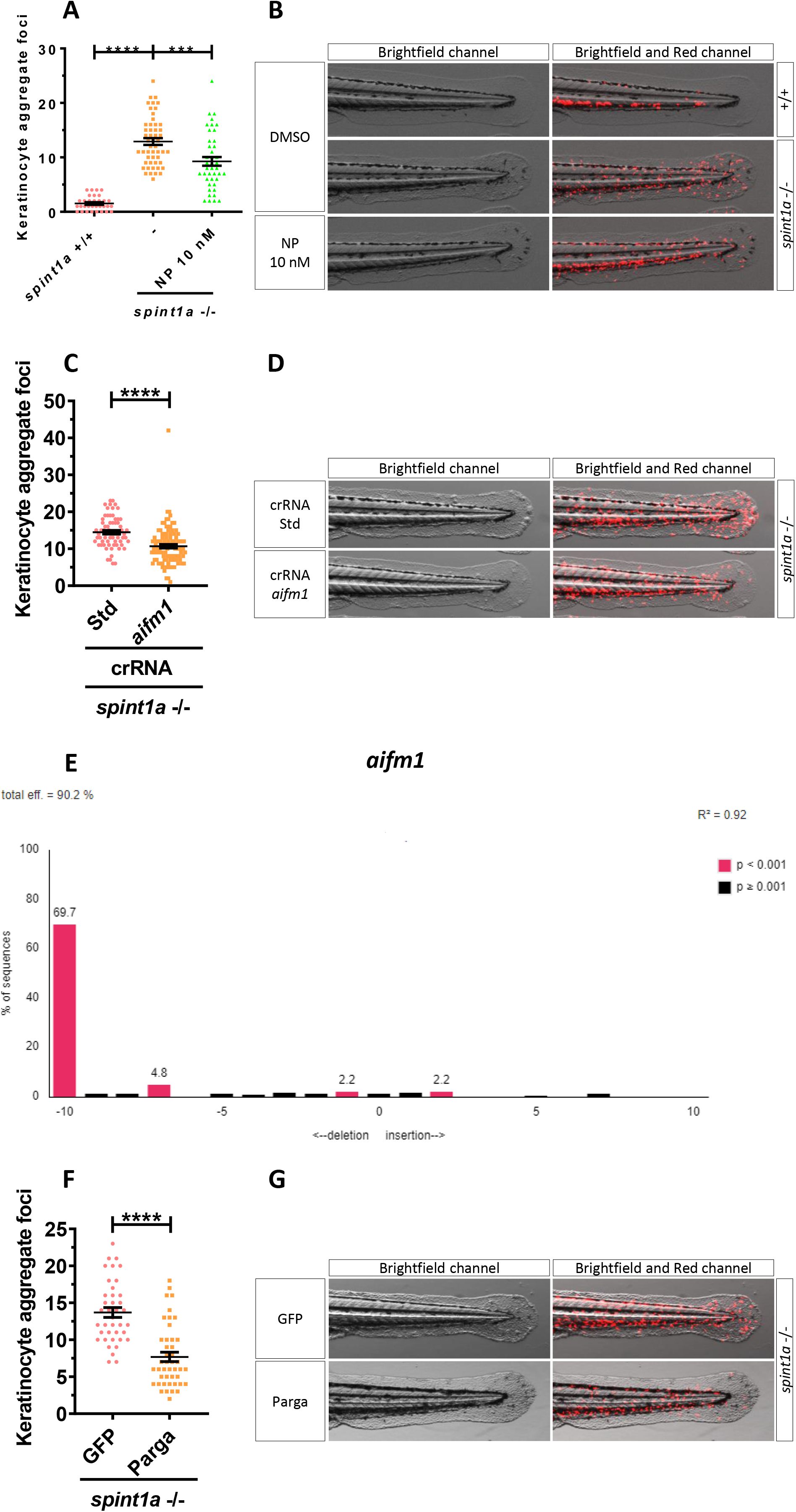
Inhibition of parthanatos rescues morphological skin alterrations of Spint1a-deficient larvae. Quantification of keratinocyte aggregates (A, C, F) and detailed representative merge images (brightfield and red channels) (B, D, G) of wild type and Spint1a-deficient larvae treated with vehicle (DMSO) or 10 nM N-phenylmaleimide (NP) (A, B), *aifm1* genetic inhibition (C, D) and *parga* mRNA overexpression (F, G) of zebrafish larvae shown in Figure 5. Analysis of genome editing efficiency or larvae injected with control or *aifm1* crRNA/Cas9 complexes and quantification rate of non-homologous end-joining mediated repair (NHEJ) showing all insertions and deletions (INDELs) (https://tide.deskgen.com/) (E).

**Figure S7, related to Figure 6.**
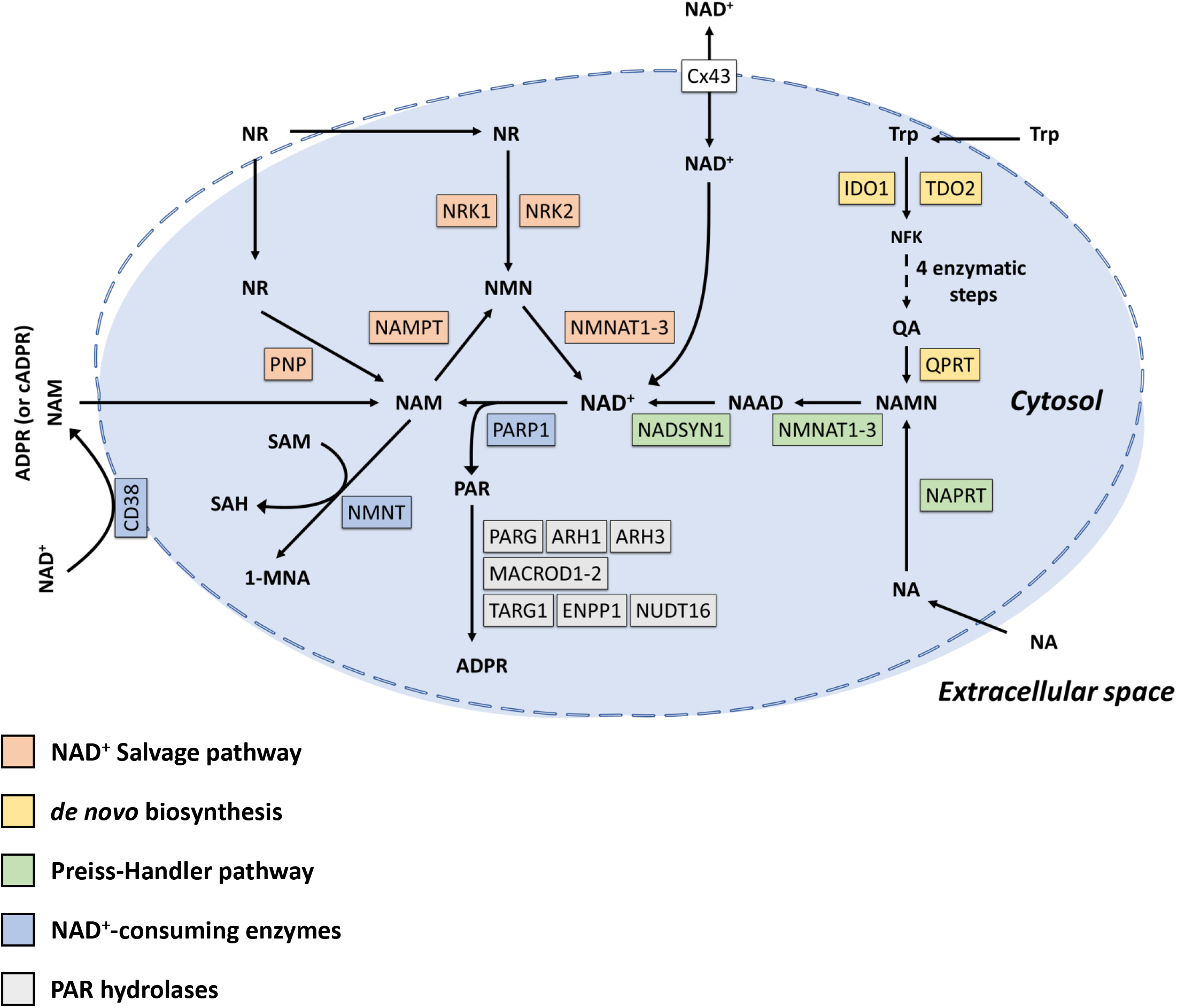
NAD^+^ and PAR metabolic pathways. NAM, NMN and NAD^+^ can be taken up by specific transporters. NAD^+^ biosynthetic pathways generate NAD^+^ from different precursors, *de novo* pathway employs dietary tryptophan (Trp) or alternatively quinolinic acid (QA), NAD^+^ Salvage pathway mainly uses nicotinamide (NAM) but nicotinamide mononucleotide (NMN) and nicotinamide riboside (NR) can also act as precursors. However, Preiss-Handler pathway utilizes nicotinic acid (NA). NAD^+^ is consumed by CD38 yielding NAM and adenosine diphosphoribose (ADPR) or cyclic ADPR (cADPR). NNMT also reduces NAD^+^ pool mediating the reaction between NAM and S-adenosylmethionine (SAM) to produce N-methylnicotinamide (1-MNA) and S-adenosylhomocysteine (SAH). Finally, PARP1 synthesizes PAR by using NAD^+^ as a cofactor. PAR is degraded to ADPR mediated by different PAR hydrolases which cleave specific chemical linkages (exo- or endoglycosidically). Metabolic intermediates: N-formylkynurenine (NFK), nicotinic acid adenine dinucleotide (NAAD) and nicotinic acid mononucleotide (NAMN). NAD^+^ transporter: connexin 43 (CX43).

**Figure S8, related to Figure 6.**
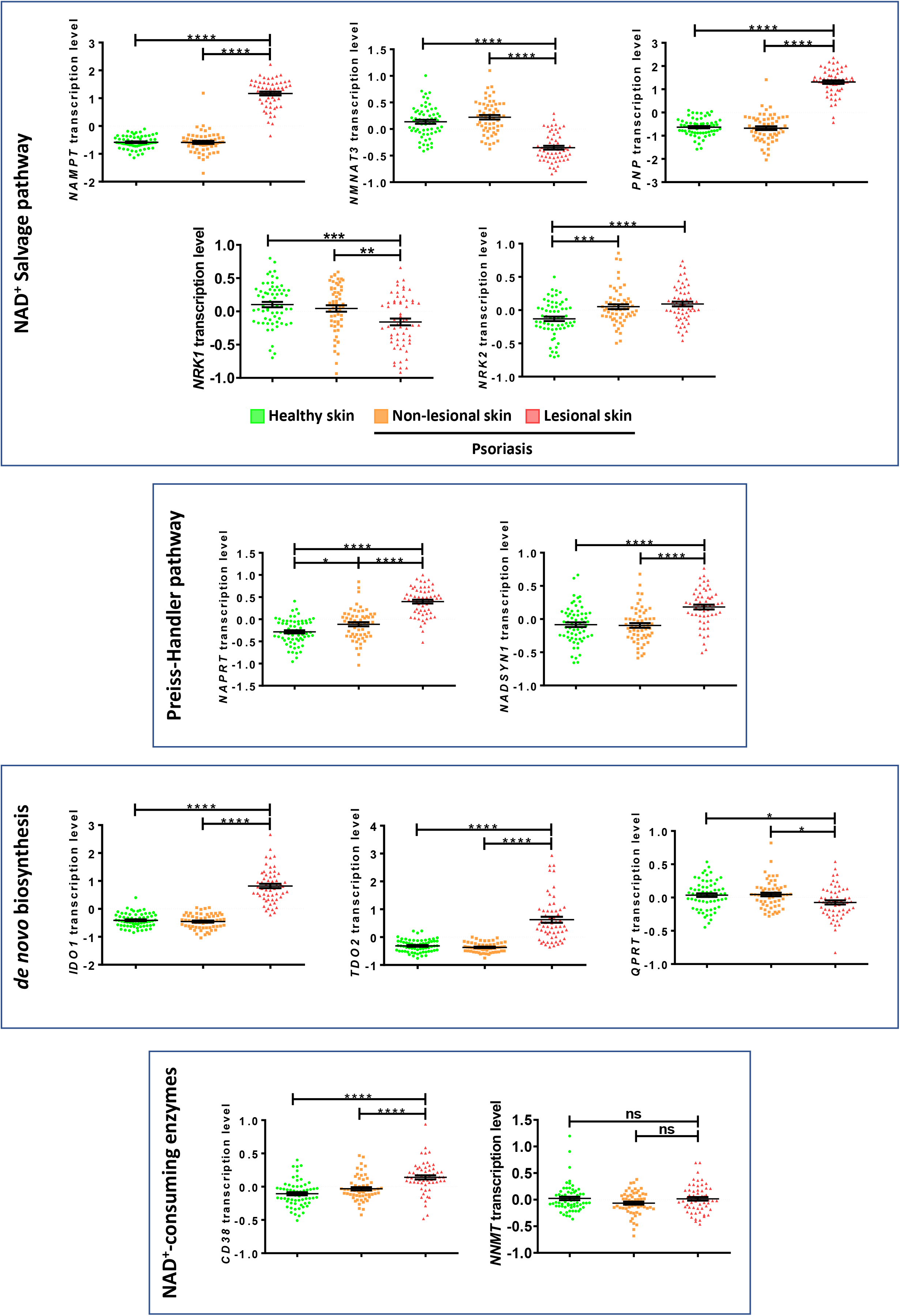
Differential expression profiles of genes encoding key NAD^+^ metabolic enzymes in psoriasis and atopic dermatitis. Transcriptomic data from human psoriasis (GDS4602) samples from the Gene Expression Omnibus (GEO) database. Non-lesional and lesional psoriasis skin were compared with heathy skin samples. Each dot represents one individual and the mean ± S.E.M. for each group is also shown. P values were calculated using one-way ANOVA and Tukey multiple range test (A) and t-Test (B). ns, not significant. *p≤0.05, **p≤0.01, ***p≤0.001, ****p≤0.0001.

**Figure S9, related to Figure 6.**
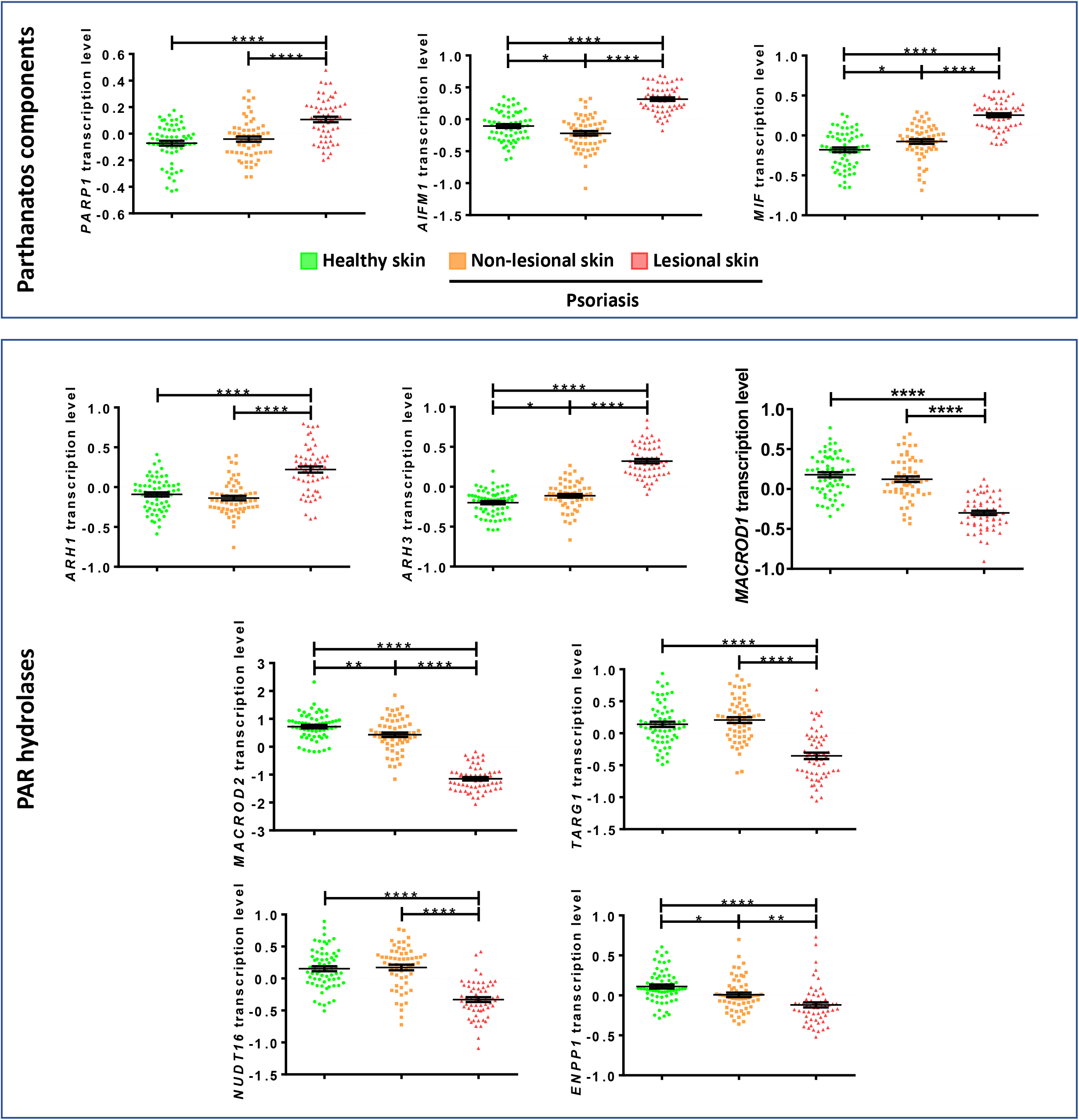
Differential expression profiles of genes encoding parthanatos components in psoriasis and atopic dermatitis. Transcriptomic data from human psoriasis (GDS4602) samples from the Gene Expression *Omnibus* (GEO) database. Non-lesional and lesional psoriasis skin were compared with heathy skin samples. Each dot represents one individual and the mean ± S.E.M. for each group is also shown. P values were calculated using one-way ANOVA and Tukey multiple range test (A) and t-Test (B). ns, not significant. *p≤0.05, **p≤0.01, ***p≤0.001, ****p≤0.0001.

**Table S1.**
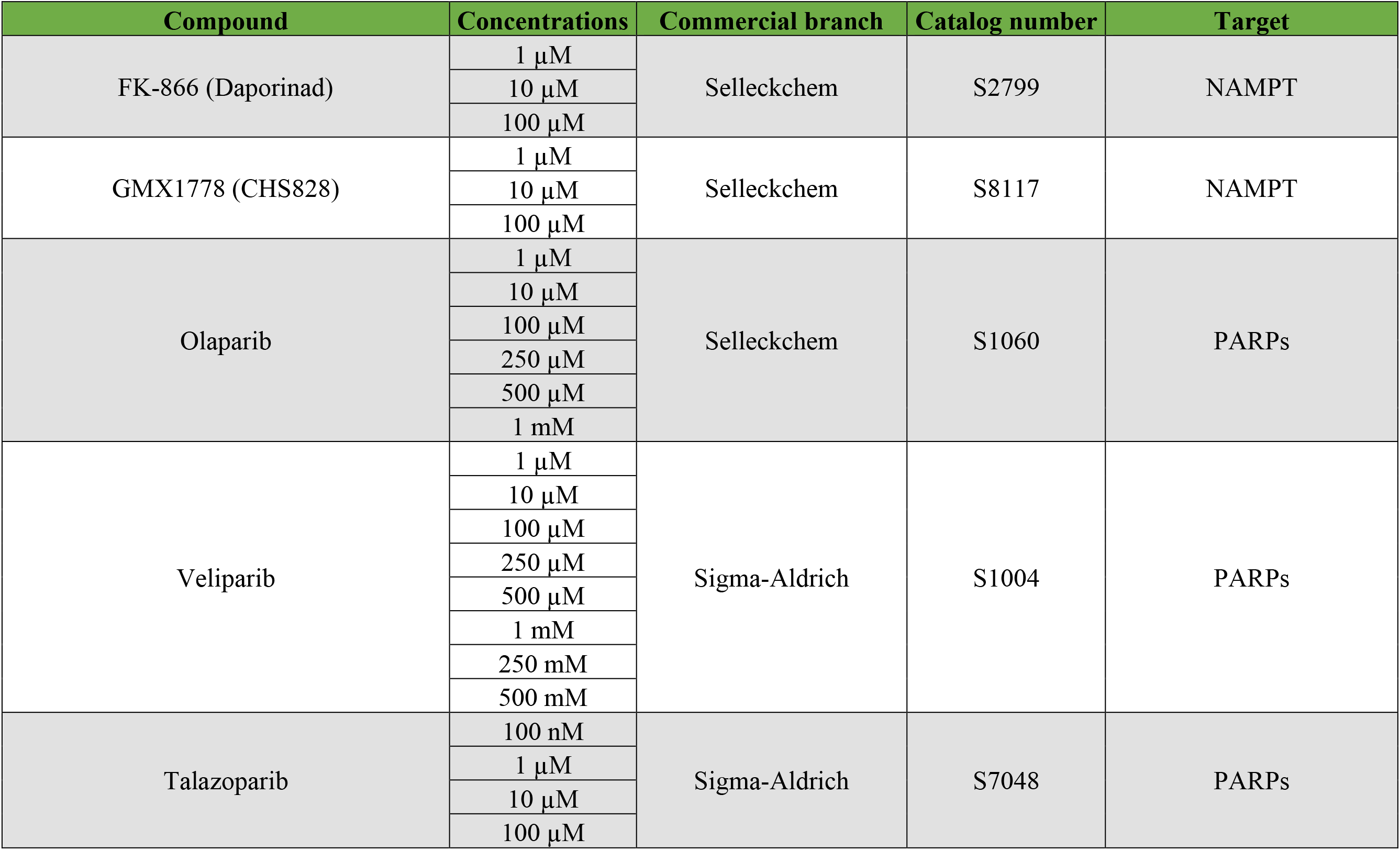
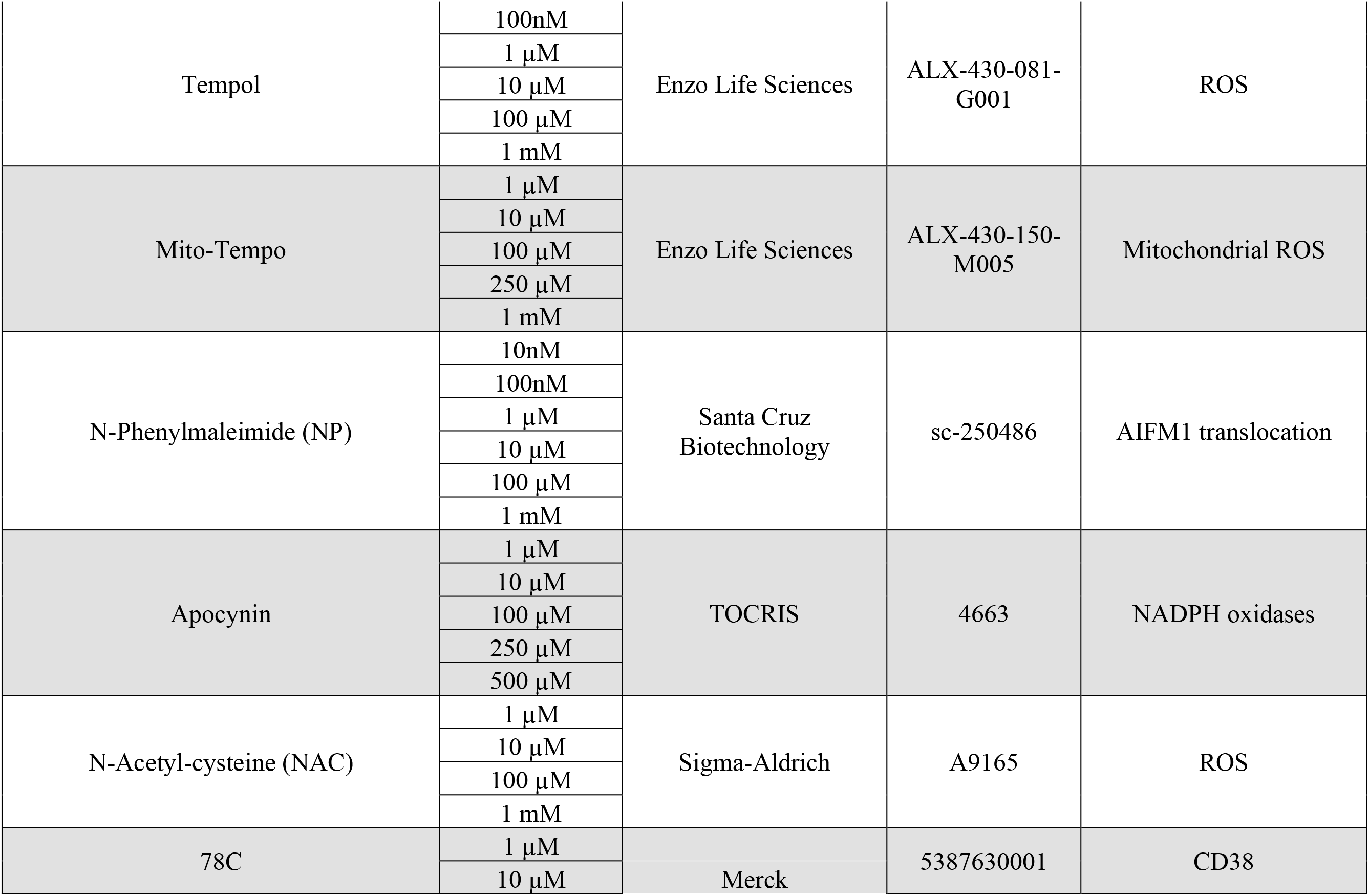
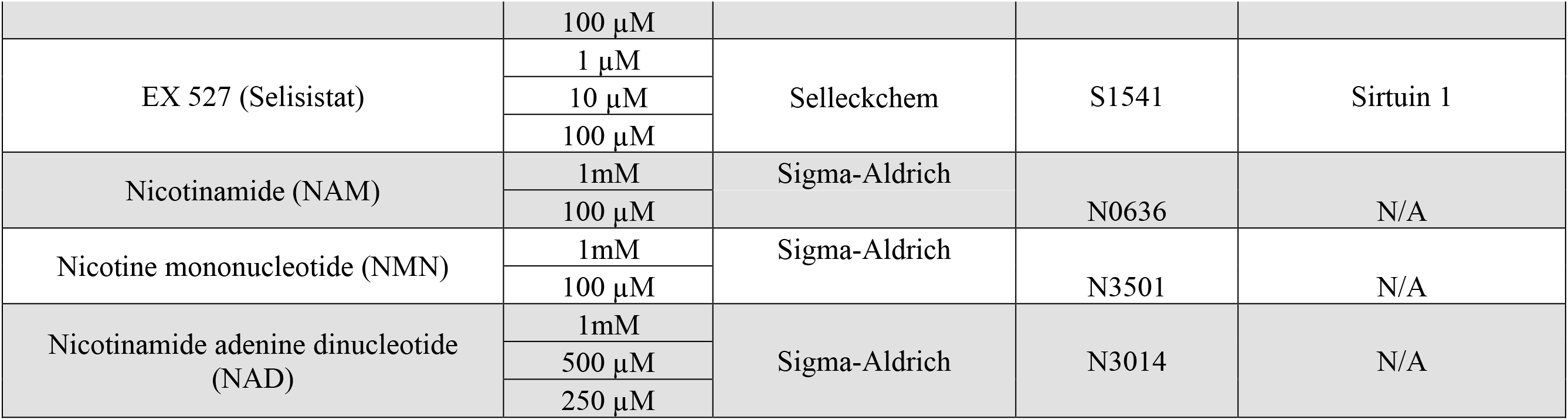
Compounds used in this study.

